# Unraveling the relation between EEG-correlates of attentional orienting and sound localization performance: a diffusion model approach

**DOI:** 10.1101/616573

**Authors:** Laura-Isabelle Klatt, Daniel Schneider, Anna-Lena Schubert, Christina Hanenberg, Jörg Lewald, Edmund Wascher, Stephan Getzmann

## Abstract

Understanding the contribution of cognitive processes and their underlying neurophysiological signals to behavioral phenomena has been a key objective in recent neuroscience research. Using a diffusion-model framework, we investigated to what extent well-established correlates of spatial attention in the electro-encephalogram contribute to behavioral performance in an auditory free-field sound-localization task. Younger and older participants were instructed to indicate the horizontal position of a pre-defined target among three simultaneously presented distractors. The central question of interest was whether posterior alpha lateralization and amplitudes of the anterior contralateral N2 subcomponent (N2ac) predict sound localization performance (accuracy, mean reaction time) and/or diffusion model parameters (drift rate, boundary separation, non-decision time). Two age groups were compared to explore whether in older adults, who struggle with multi-speaker environments, the brain-behavior relationship would differ from younger adults. Regression analyses revealed that N2ac amplitudes predicted drift rate and accuracy, whereas alpha lateralization was not related to behavioral or diffusion modeling parameters. This was true irrespective of age. The results indicate that a more efficient attentional filtering and selection of information within an auditory scene, reflected by increased N2ac amplitudes, was associated with a higher speed of information uptake (drift rate) and better localization performance (accuracy), while the underlying response criteria (threshold separation), mean reaction times, and non-decisional processes remained unaffected. The lack of a behavioral correlate of post-stimulus alpha power lateralization constrast the well-established notion that pre-stimulus alpha power reflects a functionally relevant attentional mechanism. This highlights the importance of distinguishing anticipatory from post-stimulus alpha power modulations.

## 1. Introduction

When multiple sources of acoustic information are simultaneously present, selective filtering of the available information is necessary to, for instance, focus on a talker of interest while ignoring traffic noise, music playing in the background, or other peoples’ conversations. This capacity of the human auditory system is especially astonishing, given that the incoming auditory signals often overlap in time, space, or spectral content. The behavioral effects of such selective orienting of attention in noisy, multi-speaker environments, usually referred to as ‘*cocktail-party scenarios*’ (Cherry, 1953), have been studied for decades (for review, see Bronkhorst, 2015). However, the contribution of neural signals to observable behavioral performance and its underlying cognitive processes is still poorly understood. Here, we investigated the relationship between well-established correlates of spatial attention in the electro-encephalogram (EEG) and behavioral performance in an auditory sound localization task. In particular, we specified the role of modulations in the alpha frequency band as well as an anterior contralateral N2 subcomponent (N2ac; Gamble & Luck, 2011) with respect to sound-localization performance.

Lateralized modulations of alpha power amplitude have been shown to reflect the orienting of spatial attention in visual (Foster, Sutterer, Serences, Vogel, & Awh, 2017; Ikkai, Dandekar, & Curtis, 2016; Rihs, Michel, & Thut, 2007; Worden, Foxe, Wang, & Simpson, 2000), tactile (Haegens, Handel, & Jensen, 2011; Haegens, Luther, & Jensen, 2012), and auditory space (Klatt, Getzmann, Wascher, & Schneider, 2018b; Wöstmann, Herrmann, Maess, & Obleser, 2016; Wöstmann, Vosskuhl, Obleser, & Herrmann, 2018). Typically, alpha power is shown to decrease contralaterally to the attended location (Kelly, Gomez-Ramirez, & Foxe, 2009; Sauseng et al., 2005) or to increase contralaterally to the unattended or ignored location (Kelly, Lalor, Reilly, & Foxe, 2006; Worden et al., 2000). Consistently across different modalities, this lateralized pattern of alpha-band activity has been shown to be linked to visual detection performance (Händel, Haarmeier, & Jensen, 2011; Thut, Brandt, & Pascual-Leone, 2006; van Dijk, Schoffelen, Oostenveld, & Jensen, 2008), tactile discrimination acuity (Craddock, Poliakoff, El-deredy, Klepousniotou, & Lloyd, 2017; Haegens et al., 2011), and listening performance (Tune, Wöstmann, & Obleser, 2018; Wöstmann et al., 2016). Going beyond a mere correlational approach, recent studies applying stimulation techniques, such as transcranial magnetic stimulation (TMS) or continuous transcranial alternating current stimulation (tACS), suggest a causal role of alpha oscillations in the processing of incoming information (Romei, Gross, & Thut, 2010; Wöstmann et al., 2018). Two major (not necessarily mutually exclusive) mechanisms have been proposed to underlie those asymmetric modulations of alpha power oscillations: target enhancement (Noonan et al., 2016; Yamagishi, Goda, Callan, Anderson, & Kawato, 2005) and distractor inhibition (Kelly, Lalor, Reilly, & Foxe, 2006; Rihs et al., 2007; Schneider, Göddertz, Haase, Hickey, & Wascher, 2019; Worden et al., 2000). While the majority of previous studies investigated pre-stimulus alpha oscillations as an index of anticipatory allocation of spatial attention in young adults, we focused on post-stimulus alpha lateralization in a sound-localization task, simulating a ‘*cocktail-party scenario*’. Such an experimental setup more closely resembles frequent real-life situations, in which a person searches for a sound of interest (e.g., a voice or a ringing phone) without knowing in advance where to look for it. In fact, there is first evidence that distinct attentional mechanisms contribute to the preparation for as opposed to the ongoing processing of a stimulus (van Ede, Szebényi, & Maris, 2014). In addition, we explore whether the proposed mechanistic function of alpha oscillations extends to samples of older participants, which remains an ongoing matter of debate (Hong, Sun, Bengson, Mangun, & Tong, 2015; Mok, Myers, Wallis, & Nobre, 2016; Tune et al., 2018; Vaden, Hutcheson, McCollum, Kentros, & Visscher, 2012).

A second neural measure of interest, indicating the allocation of attention within an auditory scene, is the N2ac. The N2ac has been shown to be evoked in the N2 latency range (starting at around 200 ms) when detecting or localizing a target sound in the presence of one or multiple distractor stimuli, using artificial sounds (Gamble & Luck, 2011), animal vocalizations (Klatt, Getzmann, Wascher, & Schneider, 2018a; Lewald & Getzmann, 2015), or spoken numerals (Lewald, Hanenberg, & Getzmann, 2016). While the N2ac was originally suggested to reflect the allocation of selective attention to the target (Gamble & Luck, 2011), analogously to the visual posterior contralateral N2 subcomponent (N2pc; Eimer, 1996; Luck & Hillyard, 1994), its functional significance remains ambiguous. Here, we aimed to provide further evidence on the functional significance of the N2ac by investigating its relationship to sound localization performance.

In the present study, the diffusion modelling approach (Ratcliff, 1978) was applied, allowing for a more detailed understanding of behavioral patterns in discrimination tasks (for recent reviews, see Ratcliff & McKoon, 2008; Voss, Nagler, & Lerche, 2013). Although diffusion models are still only rarely used in cognitive neuroscience research (see, e.g., Nunez, Vandekerckhove, & Srinivasan, 2017; Philiastides, Ratcliff, & Sajda, 2006; Ratcliff, Philiastides, & Sajda, 2009; Schubert, Hagemann, Voss, Schankin, & Bergmann, 2015; Schubert, Nunez, Hagemann, & Vandekerckhove, 2018a), the interest in and the application of this methodological approach has increased considerably during the past decade. The general purpose of diffusion models is to decompose the cognitive processes underlying a binary decision. As one of the major advantages of the diffusion model, the estimation procession is not limited to single mean or median values, but takes the whole reaction time (RT) distribution into account. Specifically, the resulting separation of processing components offers an enormous potential to provide more detailed descriptions of cognitive processes and to generate more accurate predictions for behavioral and neurophysiological data (Ratcliff & McKoon, 2008; Turner, Rodriguez, Norcia, McClure, & Steyvers, 2016).

The diffusion model assumes that in order for a decision to be made and a reaction to be executed, evidence for either response is accumulated in the course of a noisy process until it reaches either the decision boundary of response A or response B (see figure 2 in Voss et al., 2013 for an illustration of this evidence accumulation process). The basic diffusion model includes the following parameters: The drift rate *v* describes the speed at which evidence is accumulated (or “the rate of accumulation of information”, Ratcliff & McKoon, 2008, p. 3), with higher drift rates resulting in shorter RTs and fewer errors. Threshold separation *a* indicates the amount of information considered until a decision is made. That is, conservative response criteria that are associated with slower but more accurate responses, result in large estimates of *a*, while more liberal response criteria result in smaller estimates of *a*. Threshold separation and drift rate have been shown to be negatively correlated, due to the fact that individuals with higher drift rates tend to allow more liberal response criteria (i.e., smaller threshold separation values; Schmiedek, Oberauer, Wilhelm, Süß, & Wittmann, 2007). A priori biases towards one of the decision thresholds are reflected by starting point *z*. Beyond that, the model also includes non-decisional processing, such as response execution, working memory access, or stimulus encoding. The latter is indicated by the non-decision time constant t_0_. Typically, older adults show a slowing in this decision-unrelated domain (Ratcliff, Thapar, & McKoon, 2001; Thapar, Ratcliff, & McKoon, 2003). Finally, trial-to-trial variability in drift rate (s_v_), non-decision time (s_t0_), starting point (s_z_), and the proportion of contaminated trials (*p*_*diff*_; e.g., underlying non-diffusion like processes) can be accounted for.

In summary, here we aimed at characterizing the relation between electrophysiological correlates of attentional orienting within a complex auditory scene (i.e., alpha lateralization and N2ac) and sound-localization performance, which was assessed by classical RT and accuracy measures as well as by diffusion modeling parameters. We hypothesized that, if the cognitive processes reflected by alpha power modulations and N2ac amplitudes contribute to the successful selection of the target from a sound array containing simultaneously present distractors, they should in turn contribute to the information accumulation process that results in the localization of the target. Hence, alpha power modulations and N2ac amplitudes should predict drift rate (i.e., the speed of information accumulation), and in turn, RT and accuracy.

The data analyzed here were taken from a separate study on effects of auditory training on cocktail-party listening performance in younger and older adults (Hanenberg, Getzmann, & Lewald, unpublished). Exclusively pre-training data of this study were used. The sample analyzed here included both age groups. Although we did not primarily aim at the investigation of age effects, age differences with respect to sound localization performance, alpha lateralization and N2ac, as well as the relation between these electrophysiological correlates of attentional orienting and sound-localization performance were considered. Irrespective of the expected age-related decline, we proposed the latter brain-behavior relationship to be true for both age groups.

## 2. Materials and methods

### 2.1 Participants

The original sample included 28 older adults and 24 younger adults. Data for three younger participants were discarded because of technical problems with the EEG recording. In addition, two older participants were excluded from analysis since their performance was below (14% correct) or very close to (30%) chance level (25%). Consequently, the final sample included 26 older adults (mean age 69 yrs, range 56–76 yrs, 13 female) and 21 younger adults (mean age 24 yrs, range 19–29 yrs, 11 female). All participants were right-handed as assessed by the Edinburgh Handedness Inventory (Oldfield, 1971).

An audiometry, including eleven pure-tone frequencies (0.125 - 8 kHz; Oscilla USB100, Inmedico, Lystrup, Denmark) was conducted. Hearing thresholds in the speech frequency range (< 4kHz) indicated normal hearing (≤ 25 dB) for all younger participants and mild impairments for older participants (≤ 40 dB). The study was conducted in accordance with the Declaration of Helsinki and was approved by the Ethical Committee of the Leibniz Research Centre for Working Environment and Human Factors. All participants gave their written informed consent for participation.

### 2.2 Experimental setup, procedure, and stimuli

The original study, in which data were collected (Hanenberg, Getzmann, & Lewald, unpublished), comprised three training sessions on three days, with three experimental blocks per session (15 min pre-training; 15 min post-training; 1 h post-training) and with intervals of 1-3 weeks between sessions. For the present re-analysis, exclusively the data obtained in the pre-training blocks, pooled across the three sessions, were used. The experiment was conducted in a dimly lit, echo-reduced, sound-proof room. Participants were seated in a comfortable chair that was positioned with equal distances to the left, right, and front wall of the room. Participants’ head position was stabilized by a chin rest. A semicircular array of nine broad-band loudspeakers (SC5.9; Visaton, Haan, Germany; housing volume 340 cm^3^) was mounted in front of the participant at a distance of 1.5 meters from the participant’s head. Only four loudspeakers, located at azimuthal positions of −60°, −20°, 20°, and 60°, were used for the experimental setup of the present study. A red light-emitting diode (LED; diameter 3 mm, luminous intensity 0.025 mcd) was attached right below the central loudspeaker in the median plane of the participant’s head at eye level. The LED was continuously on and served as a central fixation point.

The sound localization task applied in the present study was a modification of the multiple-sources approach that has been used in several previous studies on auditory selective spatial attention in “cocktail-party scenarios” (Lewald, 2016, 2019; Lewald & Getzmann, 2015; Zündorf, Karnath, & Lewald, 2011, 2014; Zündorf, Lewald, & Karnath, 2013). Details of the present task version have been previously described in detail (Lewald et al., 2016). Briefly, participants indicated the position of a pre-defined target numeral that was presented simultaneously with three distractor numerals. The target was kept constant for each participant and was counterbalanced across participants and age groups such that each numeral served as a target an equal number of times within the overall experiment. Four one-syllable numerals (“eins”, 1; “vier”, 4; “acht”, 8; “zehn”, 10), spoken by two male (mean pitch 141 Hz) and two female (mean pitch 189 Hz) native German speakers, served as sound stimuli (Lewald et al., 2016). All numerals were presented equally often at each of the four possible loudspeaker positions (located at −60°, −20°, 20°, 60° azimuth). Numerals presented in each trial were spoken by four different speakers. The overall sound pressure level of the sound arrays was 66 dB(A), as measured at the position of the participant’s head using a sound level meter with a ½” free-field measuring microphone (Types 2226 and 4175, Brüel & Kjær, Nærum, Denmark).

The target was present in each trial, with target position, distractor positions, and speakers varying between trials following a fixed pseudo-random order. The stimulus duration was 600 ms, followed by a response period of 2 s and an inter-trial interval of 525 ms, resulting in a total trial duration of 3.125 s. The response was given by pressing one out of four response buttons with the index finger of the right hand. The response buttons were arranged in semi-circular array, related to the four possible target locations (i.e., far left, inner left, inner right, far right). Each block consisted of 288 trials, resulting in a duration of 15 minutes per block. As already mentioned above, data from three blocks, assessed on different days, were pooled. Thus, there was a total of 864 trials per participant. On each of the three days, participants completed a short training sequence of ten trials prior to the experiment to familiarize themselves with the task.

### 2.3 EEG recording and pre-processing

The continuous EEG was recorded from 58 passive Ag/AgCl electrode cap at a sampling rate of 1 kHz using a QuickAmp-72 amplifier (Brain products, Gilching, Germany). The electrode montage was arranged according to the international 10/10 system. Two electrodes were positioned on the left and right mastoid, respectively. In addition, two horizontal and two vertical electro-oculography (EOG) electrodes were placed below and above the right eye and at the outer canthi of the left and right eye, respectively. The ground electrode was positioned right above the nasion, in the center of the forehead. The average of all electrodes served as an online-reference. Electrode impedances were kept below 10 kΩ.

Offline pre-processing of the data was conducted using the open-source toolbox EEGLAB (v14.1.2b, Delorme & Makeig, 2004) for Matlab (R2018a). The continuous EEG data were high-pass filtered at 0.5 Hz (6601-point FIR filter, band width 0.5 Hz, cut-off frequency 0.25 Hz) and low-pass filtered at 30 Hz (441-point FIR filter, transition band width 7.5 Hz, cut-off frequency 33.75 Hz). Using the automated channel rejection procedure implemented in EEGLAB, channels with a normalized kurtosis greater than five standard deviations of the mean were rejected. The data were re-referenced to the average of all remaining EEG electrodes (including 2 mastoid electrodes) and segmented into epochs ranging from −1000 to 3125 ms, relative to sound array onset. For epoched data, the 200 ms time window prior to sound array onset served as a baseline. Independent component analysis (ICA) was run on a subset of the original data, downsampled to 200 Hz and containing only every second trial. The derived independent component (IC) decomposition was then projected onto the original dataset with a 1 kHz sampling rate comprising all trials. Using the DIPFIT plugin of the EEGLAB toolbox, a single-equivalent current dipole model was computed for each of the independent component scalp maps by means of a spherical head model (Kavanagh, Darcey, Lehmann, & Fender, 1978). Artefactual ICs were identified and excluded in two subsequent steps: The automated algorithm ADJUST (Mognon, Jovicich, Bruzzone, & Buiatti, 2011) was applied to identify and reject components related to blinks, eye movements, and generic discontinuities. In addition, because artefactual independent components usually do not resemble the projection of a single dipole (Onton & Makeig, 2006), all components with a residual variance exceeding 40% of the dipole solution were rejected. The resulting IC solution was visually inspected for any additional artefactual components that were not detected by this automated rejection procedure. On average, 28 independent components (out of 51 to 58 ICs) were rejected for each participant (range: 12 – 40). Finally, the automated artifact rejection procedure implemented in EEGLAB (threshold limit: 1000 μV, probability threshold: 5 standard deviations) was performed. On average, the procedure rejected 207 trials (range: 92 – 317), that is, 23% of trials (range: 10.6 – 36.6%). Only trials with correct responses were submitted to further analyses of EEG data (cf. *2.4.2* and *2.4.3*). Data from channels, which were originally rejected, were reconstructed using EEGLAB’s spherical interpolation procedure.

### 2.4 Data analyses

#### 2.4.1 Behavioral data

Behavioral performance was quantified by means of mean RTs and mean accuracy (proportion of correct trials) as well as diffusion model parameters. Proportion of correct trials included only responses that were given within the maximum response period of 2 seconds. A total of 83 trials (i.e., on average 2 trials per participant, range = 0 – 16, median = 1) were rejected as incorrect due to missing responses and responses that exceeded the maximum response period.

The diffusion modeling framework was applied to the present auditory localization task, in which participants were instructed to localize the position of a given target within a four-sound array of one-syllable spoken numerals. Because the diffusion model is originally based on a two-choice decision task, the decision process is here assumed to present a continuous accumulation of evidence for the true target location relative to the three non-target locations. Previous applications of the diffusion model have shown that it can validly describe decision making in four-choice alternative response tasks (Schubert et al., 2015; Schubert, Nunez, Hagemann, & Vandekerckhove, 2018b). To eliminate outliers that could bias model results (Voss, Voss, & Lerche, 2015), extremely fast (< 150 ms) and extremely slow (> 3000 ms) RTs were discarded. Subsequently, data were log-transformed and *z*-standardized in order to exclude all trials with RTs exceeding ± 3 standard deviations of the mean for each individual participant.

The free software fast-dm (Voss & Voss, 2007) was used to fit a diffusion model to the RT distributions of the present data. The model parameters were estimated based on an iterative permutation process using the Kolmogrov Smirnov test statistic. The starting point *z* was set to 0.5, presuming that participants were not biased towards one of the two response categories (correct target location versus distractor location). The parameters *a*, *v*, and t_0_ were allowed to vary freely. In addition, parameters *s*_*v*_ and *st*_*0*_ were estimated because they led to a notable improvement of model fit. Trial-to-trial variability of starting point (*s*_*z*_), the difference in speed of response execution (*d*), as well as the measure for the percentage of contaminants (*p*_*diff*_) were set to 0. To graphically evaluate model fit, we plotted observed versus predicted accuracy as well as observed versus predicted values of the RT distribution for the first (.25), second (.50), and third (.75) quartile. Predicted parameter values were derived using the *construct-samples* tool of fast-dm (Voss & Voss, 2007). That is, 500 datasets were generated for each participant based on each individual’s empirical parameter values and number of trials. Finally, the mean quartile values and mean response accuracy were calculated for each participant. Pearson correlations were calculated to quantify the relationship between empirical data and model predictions for both age groups. If the majority of data points lies close to the line of perfect correlation, good model fit can be assumed.

#### 2.4.2 ERP analysis

In order to investigate the N2ac component (Gamble & Luck, 2011), we computed the mean contralateral and ipsilateral ERP amplitude at fronto-central electrodes FC3/4 for older adults and FC5/6 for younger adults. The contralateral portion comprised the average signal at left-hemispheric electrodes in right target-trials and right-hemispheric electrodes in left-target trials, whereas the ipsilateral portion included the average signal at left-hemispheric electrodes in left-target trials and right-hemispheric electrodes in right-target trials. Mean amplitude was measured from 477 ms to 577 ms relative to sound array onset. The measurement window was based on a 100 ms time window set around the 50% fractional area latency (FAL; Hansen & Hillyard, 1980; Luck, 2014) in the grand average contralateral minus ipsilateral difference curve averaged across age groups and electrodes (50% FAL = 527 ms). To determine the FAL, the area under the difference curve was measured in a broad time window ranging from 200 to 800 ms relative to sound array onset. The latency at which this area is divided in two equal halves denotes the 50% FAL. We determined a common analysis time window for both age groups because a prior control analysis did not reveal any significant differences between the 50% FAL for younger (M = 525.86 ms) and older adults (M = 517.50 ms), Z = 0.26, *p* = .80, U3 = 0.48. The respective electrodes of interest (i.e. FC3/4 and FC5/6) were chosen to include the scalp sites with the most pronounced asymmetry (i.e. peak asymmetry in the age-specific grand average waveform) for each age group. This age-specific mean amplitude was measured in the timewindow specified above.

#### 2.4.3 Time-frequency data

To obtain time-frequency representations of the single-trial oscillatory power, we convolved the epoched, stimulus-locked EEG data with three-cycle complex Morlet wavelets. The number of cycles increased with frequency by a factor of 0.5, that is, half as fast as the number of cycles in the respective fast-fourier transformation (FFT). This resulted in 3-cycle wavelets at the lowest frequency (4 Hz) and 11.25-cycle wavelets at the highest frequency (30 Hz). To quantify asymmetries in the attentional modulation of total oscillatory power (induced + evoked activity), the alpha lateralization index (ALI) was calculated (Haegens et al., 2011; Wildegger, van Ede, Woolrich, Gillebert, & Nobre, 2017; Wöstmann et al., 2016). The latter quantifies the strength of the ipsilateral minus contralateral difference in alpha power relative to the total power across both hemispheres:

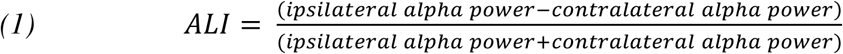

This normalization controls for potential confounds through differences in overall power level when comparing the two age groups. Mean ipsilateral and contralateral power was extracted in the alpha frequency band (8-12 Hz) at electrodes PO7/PO8 in a time window ranging from 705 to 902 ms relative to the onset of the sound array. The measurement window was based on a 200 ms time window set around the 50% FAL in the ALI difference curve averaged across age groups (50% FAL = 804 ms). The 50% FAL was calculated based on a broad time window ranging from 300 to 1400 ms relative to sound array onset. We determined a common analysis time window for both age groups, because a control analysis did not reveal any significant differences between the 50% FAL for younger (M = 796.00 ms) and older adults (M = 860.57 ms), Z = −1.27, *p* = .20, U3 = 0.31. The electrodes sites were selected based on a range of previous studies (e.g., Gould, Rushworth, & Nobre, 2011; Klatt et al., 2018; Myers, Walther, Wallis, Stokes, & Nobre, 2015; Thut, 2006; Van Der Lubbe, Bundt, & Abrahamse, 2014; van Driel, Gunseli, Meeter, & Olivers, 2017; van Ede, Niklaus, & Nobre, 2017), revealing a parieto-occipital scalp distribution and showing PO7/8 to be a representative choice of electrodes when measuring alpha lateralization. To minimize the family wise error rate, we chose to limit the analysis to one pair of electrodes. The ALI is positive when alpha power is higher over the ipsilateral hemisphere (relative to the target sound) and/or lower over the contralateral hemisphere. In contrast, negative values indicate higher alpha power contralateral to the target and/or lower alpha power over ipsilateral electrode sites. The lateralization index was calculated using the raw, baseline-uncorrected power values. ALI values for younger and older adults were submitted to parametric two-sample *t*-tests, using Satterthwaite’s approximation to assess degrees of freedom. Subsequently, one-sample *t*-tests were conducted to tests for significance of alpha lateralization within or across age groups.

#### 2.4.4 Multiple Regression

To investigate to what extent alpha lateralization and N2ac amplitudes predict behavior in the given auditory localization task, we applied regression analyses. Separate multiple linear regression models were evaluated for mean RT, drift rate *v*, threshold separation *a*, and non-decision time t_0_ as response variables, using the fitlm function implemented in the MATLAB Statistics and Machine Learning toolbox (R2018a). To account for the fact that accuracy proportions range inbetween 0 and 1, a beta regression was calculated for accuracy as a response variable using the *R betareg* package by Cribari-Neto and Zeileis (2010). For all five regression analyses, N2ac amplitudes, ALI, and age group served as predictors. In addition, to assess whether the relationship between electrophysiological correlates and behavioral outcomes differed between age groups, two interaction terms were also included (i.e., age:N2ac, age:ALI). Effects coding was used as a contrast scheme for the age group variable in order to enable a proper interpretation of lower- and higher-order effects. Model assumptions were verified by examination of residuals plots: Pearson residuals were plotted against fitted values and against predictor variables in order to assess non-constant error variance (heteroscedasticity) and deviations from linearity, respectively. In addition, normal probability plots were examined to evaluate normality of residuals. In case of a non-significant Durbin-Watson test, returning a test statistic close to 2, residuals were assumed to be uncorrelated. Variation Inflation Factors (VIFs) were inspected for signs of multicollinearity. Finally, to check for influential cases, leverage and cook’s distance were examined. Values exceeding 1 for cook’s distance (Cook & Weisberg, 1982) or 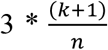 (with k indicating the number of predictors and n indicating the sample size) for leverage (Pituch & Stevens, 2016) were set as cut offs for further inspection. The inspection of residuals plots indicated deviations from normality for the drift rate regression model. Refitting the model with a log transformation (to base 10) of the drift rate values (v + 1; a constant was added to avoid negative values) resulted in approximately normally distributed residuals. Thus, ordinary least square regression was applied. For the models regarding threshold separation, non-decision time, and RT the regression model, using an iterative reweighted least squares procedure and a bisquare weight function. Adjusted R-squared (denoted as R^2^) is reported as a goodness-of-fit statistic. To correct for the fact that we conducted separate multiple regression analyses for each of the five dependent variables, *p*-values for regression coefficients were corrected using a Bonferoni-Holm procedure (Holm, 1979). Note that in each case the six p-values belonging to the same type of estimate (i.e. intercept, N2ac fixed effect, ALI fixed effect, age fixed effect, N2ac:age interaction term, or ALI:age interaction term) were corrected for multiple testing. To visualize the relationship between single predictors and outcomes, marginal effects plots (‘*ggeffect’* function from ‘ggeffects’ package, Lüdecke, 2018) and adjusted response functions (‘*plotInteraction*’ and ‘*plotAdjustedResponse*’ functions) were used for the beta regression model (in R) and linear regression models (in Matlab), respectively. Adjusted response functions describe the relationship between the fitted response and a specific predictor, while the other predictors are averaged out by averaging the fitted values over the data used in the fit. Adjusted response values are computed by adding the residual to the adjusted fitted value for each observation (Mathworks, 2019). When plotting marginal effects using ‘*ggeffect*’, the other factors are held constant at an average value (Lüdecke, 2018).

#### 2.4.5 Statistical tests and effect sizes

Data were considered normally distributed if the Lilliefors test (Lilliefors, 1967) yielded insignificant results (*p* > .05). For normally distributed data, parametric two-sample Welch’s *t*-tests were applied. Degrees of freedom were estimated using Satterthwaite’s approximation, assuming unequal variances. Wilcoxon rank-sum test served as the non-parametric counterpart in case of non-normality. To test for significance within age group, a parametric one-sample *t*-test or the non-parametric Wilcoxon signed rank test was applied. Measures of effect sizes were calculated using the MES toolbox provided by Hentschke and Stüttgen (2011). For parametric one- and two-sample *t*-tests, *g*1 and Hedge’s *g* (in the following referred to as *g*) are reported, respectively. For both measures, effect sizes of ±0.2 are typically referred to as small, values of ±0.5 as medium, and values of ±0.8 as large. For non-parametric *t*-tests, Cohen’s U3 is reported. Cohen’s U3 is a measure of overlap of two distributions, with 0.5 indicating minimal overlap and 0 or 1 indicating maximal overlap. The significance of effects was assessed at a significance level of *α* = .05. The Bonferroni-Holm correction procedure was applied to correct for multiple comparisons when appropriate (Holm, 1979). Adjusted *p*-values are denoted as *p*_adj_.

Given that *p*-values from standard inferential statistics do not allow any conclusions on whether or not the null hypothesis is true, we additionally report the Bayes factor (BF) to strengthen the interpretability of effects in the present study. In essence, the BF provides a *continuous* measure which indicates how much more likely the observed results are under a given hypothesis, compared to an alternative hypothesis (for an introduction to bayesian statistics, see Quintana & Williams, 2018; Wagenmakers et al., 2018). A BF of 1 indicates that the results are equally likely under both hypotheses (i.e. the null and the alternative hypothesis). A BF < 1 provides increasing evidence in favor of the null hypothesis relative to the alternative hypothesis, whereas a BF > 1 provides increasing evidence favoring the alternative hypothesis over the null hypothesis (Dienes, 2014). To facilitate the interpretation of BFs, the classification scheme originally proposed by Jeffreys (1961) is applied: The latter suggests that a BF > 3 and > 10 provide moderate and strong evidence for the alternative hypothesis, respectively, whereas a BF < 0.33 or < 0.1 suggests moderate and strong evidence in favor of the null hypothesis, respectively. Finally, BFs inbetween 0.33 and 3 are interpreted in terms of anecdotal evidence. However, it should be noted that those cutoffs have no absolute meaning (Dienes, 2014) in that evidence is continuous and it is directly interpretable in terms of an odds ratio (Quintana & Williams, 2018). The notation BF_10_ indicates the Bayes Factor for the alternative hypothesis (i.e., that the means of the samples are different). BF functions implemented in MATLAB by Krekelberg (2019) and the ‘BayesFactor’ package implemented in R (function: linearReg.R2stat) by Morey and Rouder (2018) were used to calculate BFs for *t*-tests and regression, respectively. In order to obtain a BF for a specific coefficient in our regression model (*BF*_*coef*_), the BF for the full model and the restricted model were compared according to the following formula: *BF*_*full*_ / *BF*_*restr*_. BF_full_ indicates the BF for the full model, including all predictors, whereas *BF*_*restr*_ indicates the BF for the restricted model, omitting the coefficient of interest. Default priors, that is, the Jeffrey-Zellner-Siow Prior for *t*-tests and a mixture of g-priors according to Liang, Paulo, Molina, Clyde, and Berger (2008) for regression, were applied. Since those packages do not support the calculation of BFs for beta regression, no bayesian statistics are provided for the regression analysis of accuracy data.

## 3. Results

### 3.1 Behavioral results

Figure 1 shows the proportion of correct responses (Figure 1a) as well as mean RTs (Figure 1b) separately for both age groups. Diffusion parameters are depicted in Figure 2. On average, younger adults showed higher accuracy (*t*(43.05) = −3.36, *p* = .002, *p*_adj_ = .01, *g* = 0.92, BF_10_ = 14.21) and faster responses than older adults (*t*(38.56) = 2.80, *p* = .008, *p*_adj_ = .038, *g* = − 0.83, BF_10_ = 6.93). The BFs indicated that the alternative model was around 14 times and 6 times more likely than the null model, respectively; thus providing strong and moderate support for a difference between age groups.

**Figure 1.**
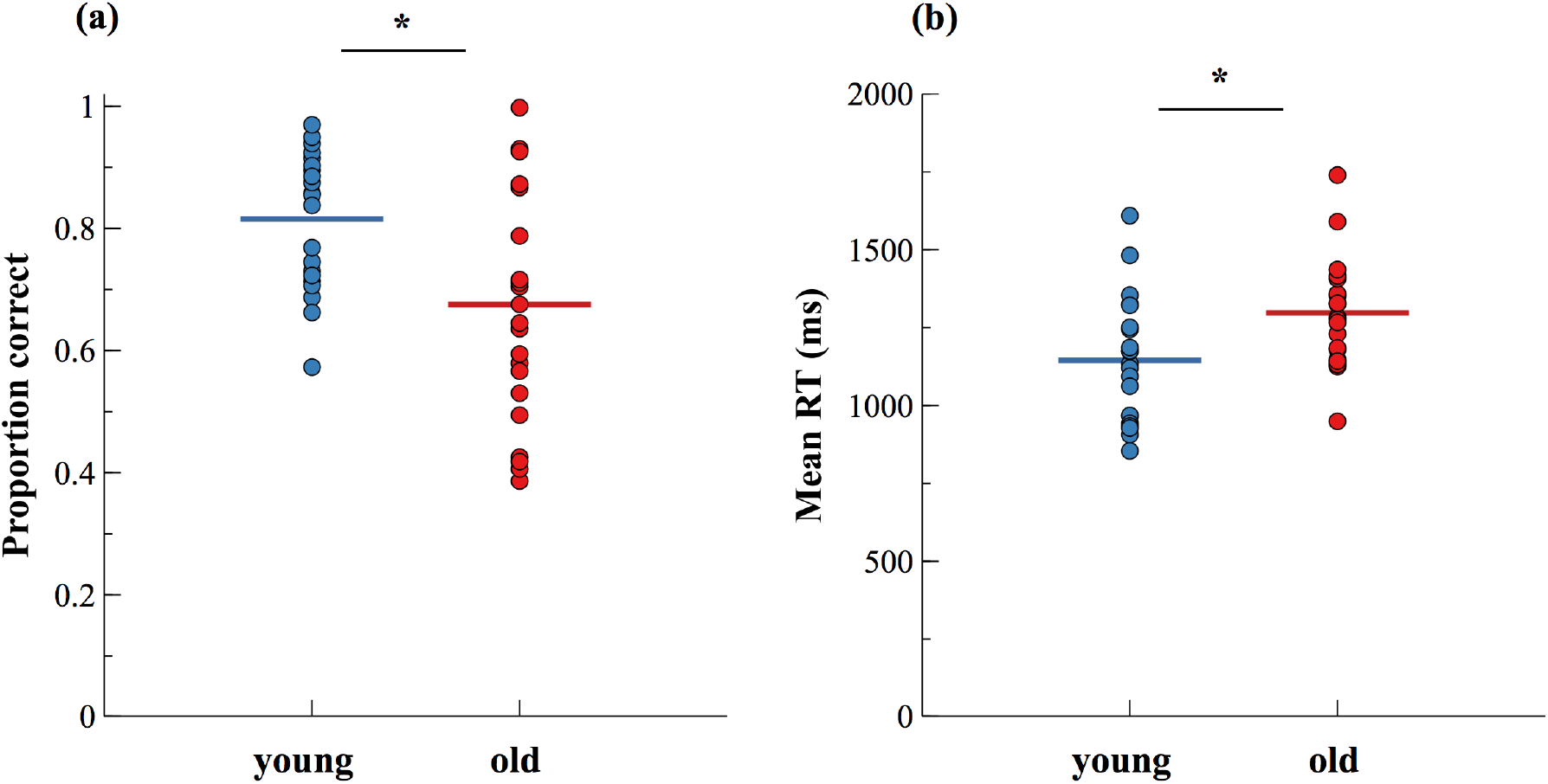
Proportion of (a) correct responses and (b) mean reaction times (RT) for younger and older adults. Colored horizontal lines indicate the respective group mean. Dots indicate individual participants’ mean values. * *p*_adj_ < .05

**Figure 2.**
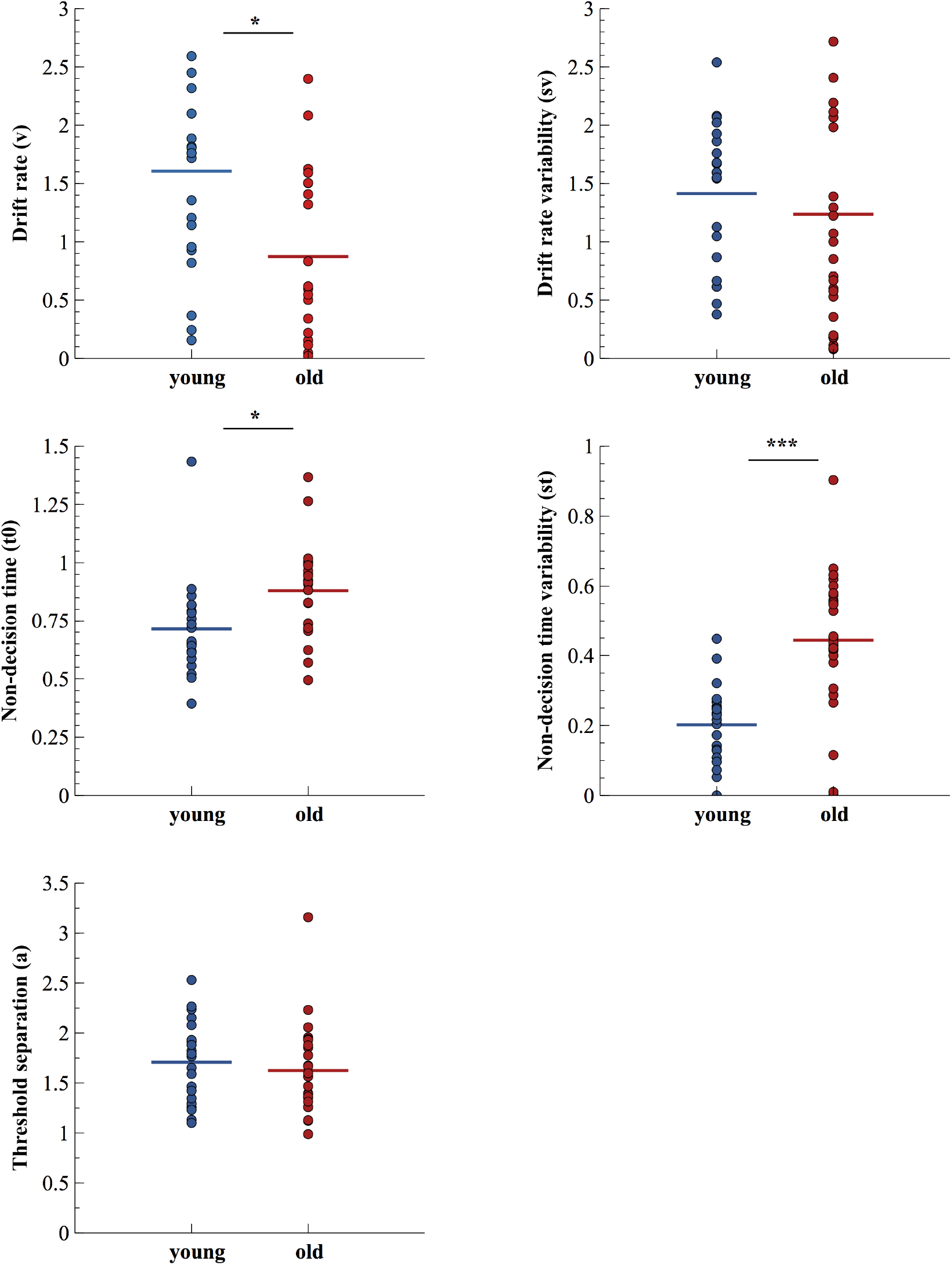
Diffusion model parameter estimates for younger and older participants. Dots represent single subject data. Colored horizontal lines show the mean model parameters within age groups. * *p*_adj_ < .05, *** *p*_adj_ < .001

While mean RTs do not offer any insights into the underlying causes of prolonged response times, diffusion parameters allow for a closer look at different possible explanations for the observed difference between age groups, including a slowdown of information update (i.e., higher drift rate *v*), a more conservative response criterion (i.e., higher threshold separation *a*), or delayed response execution (i.e., higher response constant *t*_*0*_). In our sample, older adults showed a significantly reduced drift rate (*t*_(44.89)_= −2.51, *p* = .016, *p*_adj_ = .047, g = 0.70, BF_10_ = 3.01), higher non-decision time (*t*_(41.31)_= 2.81, *p* = .008, *p*_adj_ = .038, g = −0.82, BF_10_ = 6.59) as well as higher variability of non-decision time (*t*_(40.26)_ = 5.25, *p* < .001, *p*_adj_ < .001, g = −1.43, BF_10_ = 153.9). Threshold separation values (*t*_(44.24)_ = −0.66, *p* = .513, *p*_adj_ = .513, g = 0.19, BF_10_ = 0.35) and trial-to-trial variability of drift rate (Z = −1.21, *p* = .226, *p*_adj_ = .453, U3 = 0.29, BF_10_ = 0.35) did not differ significantly between age groups. While the BFs supported classical inferential statistics for significant results (BFs > 3), they fell short of the criterion for moderate evidence for equivalence for insignificant results (BFs > 0.33). In order to graphically assess the fit of the estimated diffusion models, observed RT quartiles (.25, .5, .75) and observed accuracy were plotted against the corresponding value of the predicted distributions. As can be seen in Figure 3, the majority of data points lies close to the line of perfect correlation, indicating adequate model fit.

**Figure 3.**
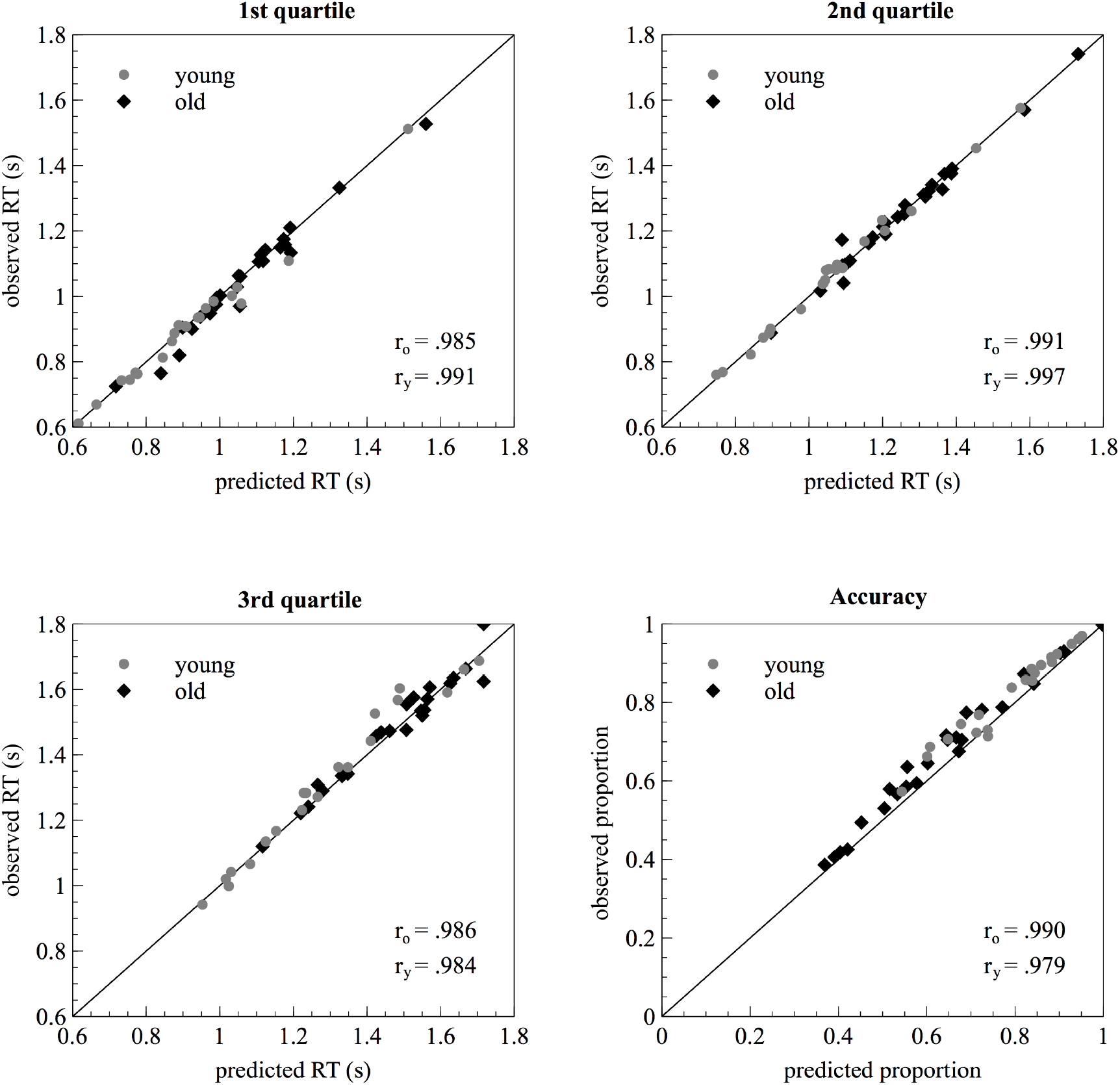
Graphical analysis of model fit. Scatter plots show the observed proportion of correct responses as well as the first three quartiles (.25, .5, .75) of the observed RT distribution as a function of the corresponding value from the predicted distribution. Dots and diamonds represent single subject data for younger and older participants.

### 3.2 N2 anterior contralateral component

Figure 4 presents the event-related potentials at fronto-central electrodes FC3/4 for older adults and electrodes FC5/6 for younger adults. In addition, the corresponding topographies based on the contralateral minus ipsilateral difference wave in the analysis time-window are depicted. N2ac amplitudes (i.e., contralateral minus ipsilateral differences) did not differ significantly between younger (M = −0.37, SD = 0.47) and older adults (M = −0.38, SD = 0.40; *t*_(39.37)_ = −0.09, *p* = .926, *g* = 0.03, BF_10_ = 0.29). The BF of 0.29 can be interpreted as insufficient evidence, supporting neither the null nor the alternative hypothesis. A one-sample *t*-test confirmed that across both age groups, N2ac amplitudes were significantly different from zero (*t*(46) = −6.02, *p* < .001, *p*_adj_ < .001, *g*_*1*_ = −0.88, BF_10_ > 1000). However, it should be noted that the original analysis time window was based on the 50% FAL in the grand average difference waveform across both age groups; thus, this procedure favors a significant result when testing overall N2ac amplitudes against zero. To avoid this problem of ‘double dipping’, we performed a second one-sample *t*-test, using a broader analysis time window of 400 to 600 ms post sound array-onset. The latter yielded comparable results (*t*_(46)_ = −4.41, *p* < .001, *p*_adj_ < .001, *g*_*1*_ = −0.64, BF_10_ > 1000). Consistently, the BF provided strong evidence in favor of the presence of an N2ac component across both age groups.

**Figure 4.**
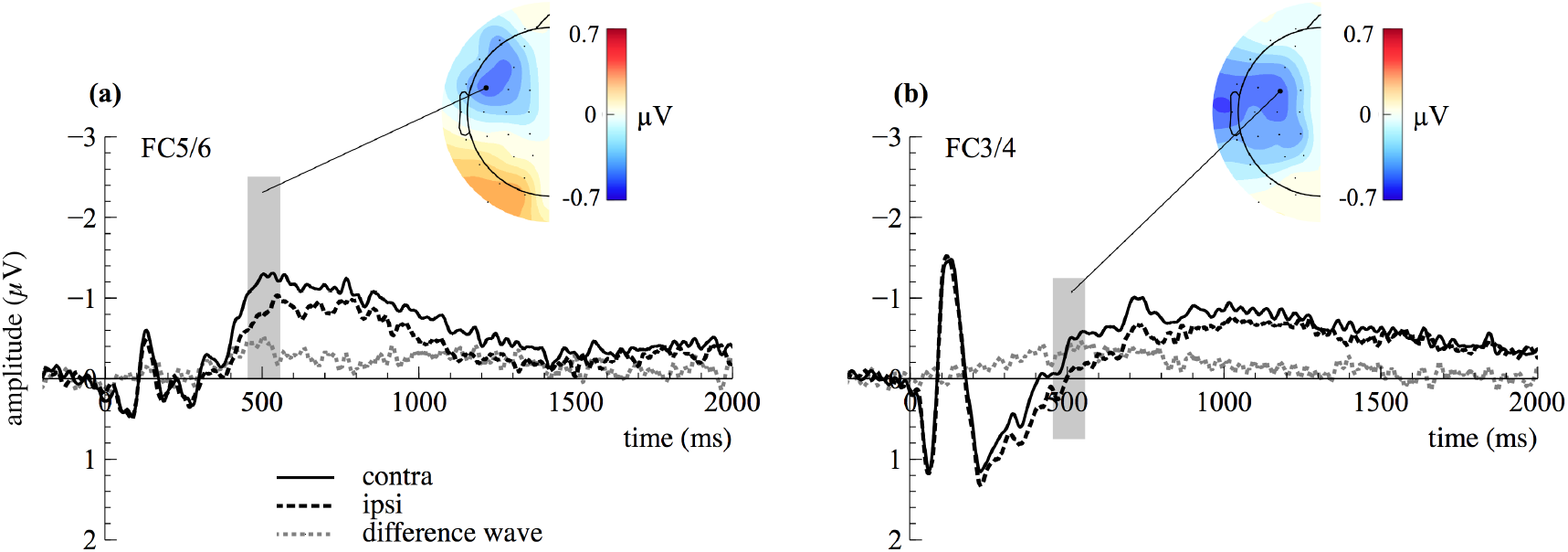
N2ac component at fronto-central electrodes FC5/6 for (a) younger and at FC3/4 for (b) older participants. Contralateral and ipsilateral portions of the signal as well as the resulting difference wave (contralateral minus ipsilateral) are depicted. Scalp topographies show the distribution of voltage differences based on the contralateral minus ipsilateral difference wave in the time window used for statistical analysis (highlighted in grey in ERP figures).

### 3.3 Alpha Lateralization

The time-frequency plot in figure 5 illustrates the asymmetric modulation of alpha power (8-12 Hz) at electrodes PO7/8 time-locked to sound-array onset, for younger (Figure 5a) and older adults (Figure 5b), respectively. In addition, the corresponding topographies based on the normalized ipsilateral minus contralateral difference in alpha power are depicted. Although younger adults appeared to show larger alpha power lateralization than older adults, the analysis revealed no significant difference in alpha power lateralization between age groups (*t*_(41.23)_ = −1.43, *p* = .161, *g* = 0.42, BF_10_ = 1.13). The BF suggested that the data were insensitive to distinguish the null (no amplitude difference between groups) from the alternative hypothesis (difference in amplitudes between age groups). Yet, a one sample *t*-test confirmed that alpha lateralization across both age groups was significantly different from zero (*t*_(46)_ = 6.07, *p* < .001, *p*_adj_ < .001, *g*_*1*_ = 0.89, BF_10_ > 1000), and the BF consistently suggested strong evidence for the alternative hypothesis. As mentioned above (cf. 3.2), the analysis time window (determined based on the 50% FAL in the grand average waveform) favors a significant result when testing across age groups, against zero. Thus, a second one-sample *t*-test was performed, based on a broader analysis time window of 600 to 900 ms post sound array onset, yielding comparable results (*t*_(46)_ = 5.91, *p* < .001, *p*_adj_ < .001, *g*_*1*_ = 0.86, BF_10_ > 1000).

**Figure 5.**
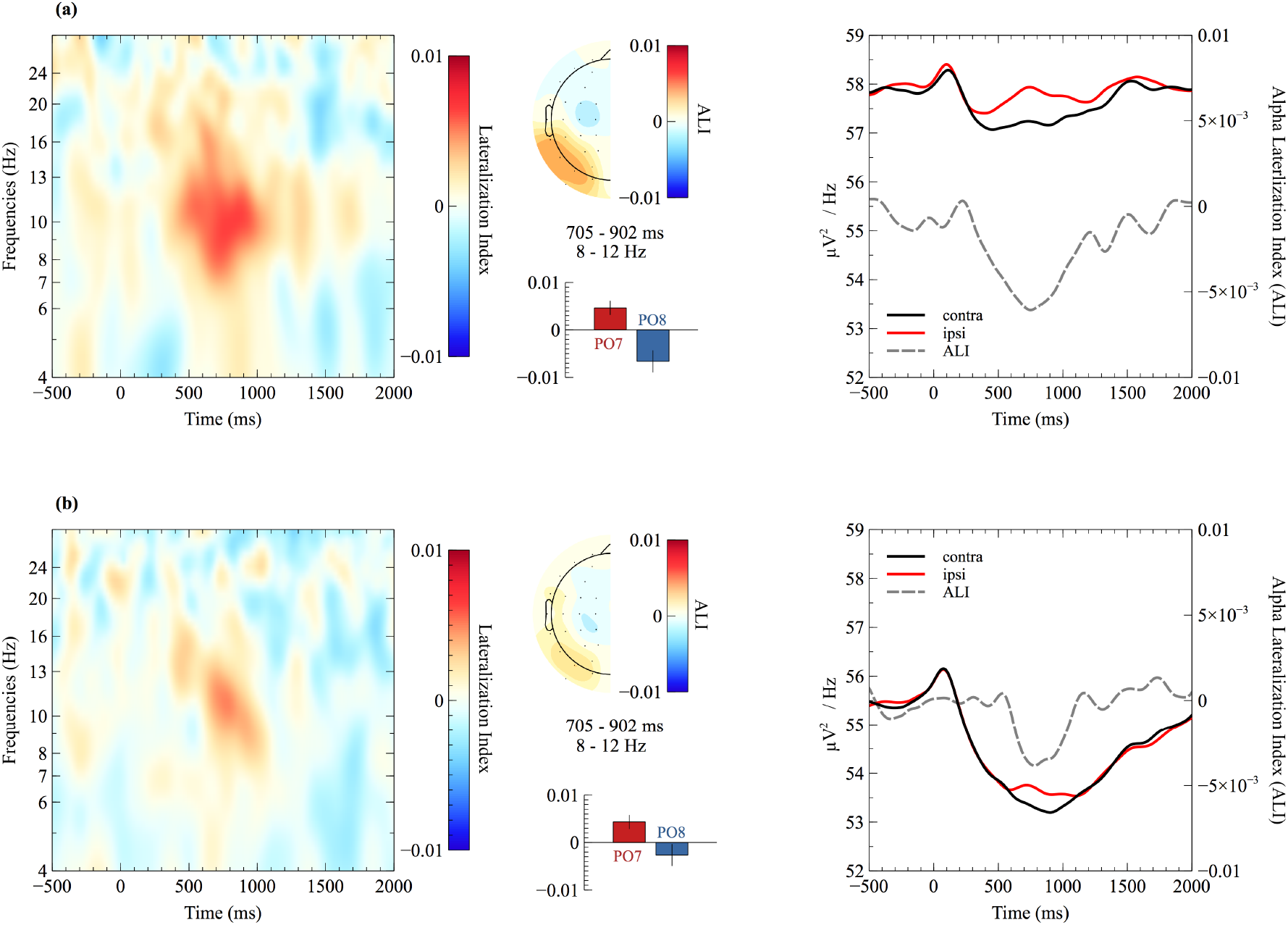
Grand average time-frequency plots of lateralization indices at electrodes PO7/8 (cf. left panel) for (a) younger and (b) older adults. The scalp topographies are based on normalized differences of ipsilateral minus contralateral alpha power in the time window used for statistical analysis. Bar graphs show the mean difference (left minus right) for the left (i.e., PO7) and right (i.e., PO8) hemisphere. Error bars indicate the standard error of the mean. Lineplots (cf. right panel) illustrate the contralateral and ipsilateral portion of the raw ERSPs as well as the resulting ALI.

### 3.4 Regression analyses

We examined the relationship between mean alpha power lateralization, N2ac amplitudes, and behavioral performance (including diffusion model parameters) using multiple linear regression. The estimated parameters are provided in Table 1. Participants with greater N2ac amplitudes showed higher accuracy (Z = −3.93, *p* < .001, *p*_adj_ < .001) and higher drift rate (*t*_(41)_ = −2.79, *p* = .008, *p*_adj_ = .032, *BF*_*coef*_ = 7.75) while there was no significant effect of alpha lateralization on those performance outcomes (accuracy: Z = −1.54, *p* = .124, *p*_adj_ = .499; drift rate: *t*_(41)_ = −0.37, *p* = .712, *p*_adj_ = 1.067, *BF*_*coef*_ = 0.43). For both models, there was no significant interaction with age (all *p*_adj_ > .160). The corresponding BFs (only available for the drift rate model, cf. 2.4.5) were below 3 (*BF*_*coef*_ ≤ 0.65) but above 0.33, thus, lending insufficient evidence for the null or the alternative hypotheses. The full models, including all predictors, explained 26% and 36% of variance in drift rate and accuracy, respectively (R^2^_adj_ = .26, F(5,41) = 4.15, *p* = .004; pseudo-R^2^ = .36, precision parameter phi = 9.73, SE = 1.96, z = 4.97, Pr(>|z|) < .001). For all other models tested, neither N2ac amplitudes nor alpha power lateralization or their interaction with age groups served as statistically significant predictors (all *p*_adj_ > .095, cf. table 1). For all but one parameter, the corresponding BFs were inconclusive (3 < *BF*_*coef*_ > 0.33), providing no substantial support for the alternative hypothesis, but neither for the null hypothesis. However, for the regression model predicting non-decision time, the BF for the interaction term N2ac*age (*p* = .095) lend moderate evidence in favor of the alternative hypothesis (*BF*_*coef*_ = 5.92), suggesting that in older adults, less pronounced N2ac amplitudes were associated with higher non-decision times. In contrast, the latter relationship appeared absent in younger adults. Age group, not surprisingly, significantly predicted non-decision times (*t*_(41)_ = 3.00, *p* = .005, *p*_adj_ = .018, *BF*_*coef*_ = 15.27), accuracy (Z = 3.03, *p* = .002, *p*_adj_ = .012), and drift rate (*t*_(41)_ = −2.86, *p* = .007, *p*_adj_ = .020, *BF*_*coef*_ = 8.78). Although age group failed to serve as a significant predictor for reaction time in the regression model framework (*t*_(41)_ = 1.82, *p* = .075, *p*_adj_ = .151, *BF*_*coef*_ = 2.38), the results largely confirm the behavioral age differences reported in section 3.1. The BF of 2.38 suggests that the data may simply be underpowered to reveal a relation between RT and age group in the present regression model. Figure 6 visualizes the reported results for those outcomes that were significantly predicted by N2ac amplitudes.

**Table 1.** Estimated parameters, standard errors, confidence intervals, and t-test (or *z*-test) statistics for each predictor in the linear (or beta) regression model.

| Outcome | v |  | a |  | t<sub>0</sub> |  | Accuracy |  | RT |  |
| --- | --- | --- | --- | --- | --- | --- | --- | --- | --- | --- |
|  | **b (SE)**<br>**[95% CI]** | *t* | **b (SE)**<br>**[95% CI]** | *t* | **b (SE)**<br>**[95% CI]** | *t* | **b (SE)** | *z* | **b (SE)**<br>**[95% CI]** | *t* |
| Intercept | 0.22***<br>(0.05)<br>[0.12 0.31] | 4.43<br>*p* < .001<br>*p*<sub>adj</sub> < .001 | 1.55***<br>(0.09)<br>[1.36 1.74] | 16.47<br>*p* < .001<br>*p*<sub>adj</sub> < .001 | 0.77***<br>(0.04)<br>[0.70 0.86] | 19.77<br>*p* < .001<br>*p*<sub>adj</sub> < .001 | 0.89***<br>(0.16) | 5.63<br>*p* < .001<br>*p*<sub>adj</sub> < .001 | 1.24***<br>(0.04)<br>[1.15 1.33] | 28.95<br>*p* < .001<br>*p*<sub>adj</sub> < .001 |
| N2ac | -0.22*<br>(0.08)<br>[-0.38 -0.06] | -2.79<br>*p* = .008<br>*p*<sub>adj</sub> = .032 | -0.32<br>(0.15)<br>[-0.62 -0.01] | -2.10<br>*p* = .042<br>*p*<sub>adj</sub> = .126 | 0.08<br>(0.06)<br>[-0.05 0.21] | 1.29<br>*p* = 0.203<br>*p*<sub>adj</sub> = .407 | -1.02***<br>(0.26) | -3.93<br>*p* < .001<br>*p*<sub>adj</sub> < .001 | 0.08<br>(0.07)<br>[-0.06 0.22] | 1.14<br>*p* = .261<br>*p*<sub>adj</sub> = .407 |
| ALI | -2.52<br>(6.77)<br>[-16.20 11.16] | -0.37<br>*p* = .712<br>*p*<sub>adj</sub> = 1.067 | -10.87<br>(13.05)<br>[-37.22 15.49] | -0.83<br>*p* = .410<br>*p*<sub>adj</sub> = 1.230 | 10.18<br>(5.45)<br>[-0.83 21.19] | 1.87<br>*p* = .069<br>*p*<sub>adj</sub> = .345 | -33.48<br>(21.81) | -1.54<br>*p* = .124<br>*p*<sub>adj</sub> = .499 | 3.73<br>(5.94)<br>[-8.27 15.74] | 0.63<br>*p* = .533<br>*p*<sub>adj</sub> = 1.230 |
| age | -0.14*<br>(0.05)<br>[-0.24 -0.04] | -2.86<br>*p* = .007<br>*p*<sub>adj</sub> = .020 | -0.07<br>(0.09)<br>[-0.26 0.12] | -0.76<br>*p* = .452<br>*p*<sub>adj</sub> = .452 | 0.12*<br>(0.04)<br>[0.03 0.20] | 3.00<br>*p* = .005<br>*p*<sub>adj</sub> = .018 | -0.48*<br>(0.16) | 3.03<br>*p* = .002<br>*p*<sub>adj</sub> = .012 | 0.08<br>(0.04)<br>[-0.01 0.16] | 1.82<br>*p* = .075<br>*p*<sub>adj</sub> = .151 |
| N2ac*age | -0.16<br>(0.08)<br>[-0.32 0.00] | -2.02<br>*p* = .050<br>*p*<sub>adj</sub> = .160 | -0.11<br>(0.15)<br>[-0.41 0.20] | -0.72<br>*p* = .474<br>*p*<sub>adj</sub> = .474 | 0.15<br>(0.06)<br>[0.03 0.28] | 2.44<br>*p* = .018<br>*p*<sub>adj</sub> = .095 | -0.53<br>(0.26) | -2.05<br>*p* = .039<br>*p*<sub>adj</sub> = .160 | 0.09<br>(0.04)<br>[-0.04 0.24] | 1.43<br>*p* = .160<br>*p*<sub>adj</sub> = .319 |
| ALI*age | -4.67<br>(6.78)<br>[-18.35 9.02] | -0.69<br>*p* = .495<br>*p*<sub>adj</sub> = .965 | -10.06<br>(13.05)<br>[-36.41 16.30] | -0.77<br>*p* = .445<br>*p*<sub>adj</sub> = 1.336 | 11.23<br>(5.45)<br>[0.22 22.24] | 2.06<br>*p* = .046<br>*p*<sub>adj</sub> = .229 | -15.30<br>(21.78) | -0.70<br>*p* = .483<br>*p*<sub>adj</sub> = 1.336 | 9.19<br>(5.94)<br>[-2.81 21.19] | 1.55<br>*p* = .130<br>*p*<sub>adj</sub> = .519 |
| Adjusted /<br>Pseudo - R<sup>2</sup> | .26 |  | .03 |  | .32 |  | .36 |  | .14 |  |
| *F*-statistic | *F*(5,41) = 4.15, *p* = .004 |  | *F*(5,41) = 1.25, *p* = .302 |  | *F*(5,41) = 5.29, *p* = .001 |  | - |  | *F*(5,41) = 2.46, *p* = .048 |  |

**Figure 6.**
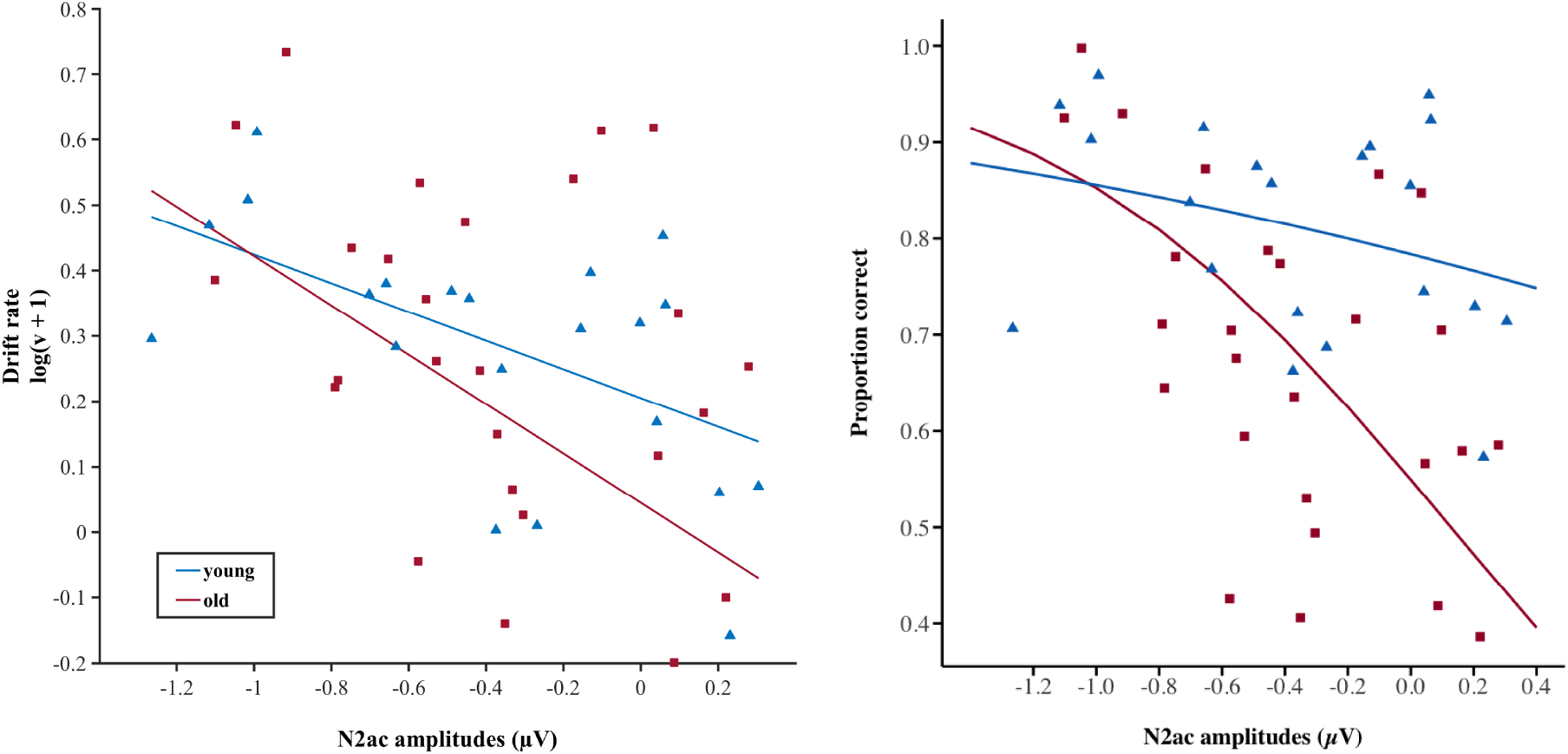
Participants’ (a) drift rate and (b) mean accuracy as a function of mean N2ac amplitude. Triangles represent younger participants (n = 21), squares represent older participants (n = 26). For the linear drift rate regression model, an adjusted response function describes the relationship between the fitted response and N2ac amplitudes, while the other predictors are averaged out by averaging the fitted values over the data used in the fit. Adjusted response data points are computed by adding the residual to the adjusted fitted value for each observation. For the accuracy beta regression model, the marginal effect of the interaction N2ac amplitude by Age group is displayed, holding the other factors constant at an average vaule.

## 4. Discussion

In the present study, we investigated the contribution of post-stimulus alpha power lateralization and N2ac amplitudes to sound localization performance in a sample of younger and older adults. Both measures have been associated with the deployment of attention in auditory space. We hypothesized that if the cortical processes reflected by alpha lateralization and N2ac amplitudes contribute to successful target selection, their magnitudes should be related to the information accumulation process (i.e., drift rate; cf. diffusion model framework in *2.5.2*), and in turn to localization accuracy and RTs. In fact, what we found only partially confirmed this hypothesis: N2ac amplitudes significantly predicted both drift rate and accuracy, while alpha lateralization was not associated with any of the behavioral outcomes. We thus proposed that N2ac and alpha lateralization reflect distinct aspects of attentional orienting in auditory scenes. Classical frequentist inferential statistics suggested that the observed relationship did not depend on age and that both age groups showed comparable neural signatures. However, bayesian alternatives to classical hypotheses testing raised doubts about these claims, suggesting that the data is inconclusive with respect to age effects in the electrophysiological data. Age differences in behavioral performance are briefly reviewed below.

### 4.1 Cocktail-party sound localization in older and younger adults

As expected, older adults showed fewer correct responses and slower response times than younger adults. This is in line with the often-described difficulties of older people to follow a conversation in noisy (“cocktail-party”) environments, which depends on the integrity of both sensory and cognitive functions (Shinn-Cunningham, 2017). Declined performance in older adults in the present task is likely to be related to age-related deficits in concurrent sound segregation (Alain & McDonald, 2007; Hanenberg, Getzmann, & Lewald, 2019; Snyder & Alain, 2005). Traditionally, such deficits have been interpreted as a result of a general sensory-cognitive decline (e.g., Pichora-Fuller et al., 2017), assuming all aspects of processing in an experimental task to be globally slowed in aging adults (Myerson, Hale, Wagstaff, Poon, & Smith, 1990). The diffusion model allows to differentiate between different aspects of processing that might be affected by age (Ratcliff, Spieler, & Mckoon, 2000): Consistent with previous results (Ratcliff et al., 2001; Ratcliff, Thapar, & McKoon, 2003, 2011), we found an increase in non-decision time for older adults. In addition, older participants varied more strongly in their non-decision time from trial to trial, indicating that this process was noisier in older adults (Spaniol, Madden, & Voss, 2006). However, rather untypically, the two age groups did not differ in their threshold separation values. This contradicts the wide-spread assumption that older adults usually aim to minimize errors (leading to more conservative response criteria), while younger adults focus on balancing speed and accuracy (Starns & Ratcliff, 2010). The observed lack of differences in response criteria between older and younger adults could be due to the relatively long response period in the present study, potentially inducing a change in task goals in younger adults. Alternatively, as the corresponding BFs were rather inconclusive, we cannot exclude that the data are simply underpowered and therefore fail to reveal significant differences in our sample. Furthermore, supporting a line of evidence that showed differences in the rate of information accumulation in some contexts (Ratcliff et al., 2011; Spaniol et al., 2006; Thapar et al., 2003), older adults had significantly decreased drift rates. Given the current state of research, the conditions under which drift rate decreases with age are still hard to grasp. Here, drift rate was significantly predicted by N2ac amplitudes. In participants with higher N2ac amplitudes (i.e., more negative difference waves) drift rates were higher, while participants with lower N2ac amplitudes tended to have lower drift rates. Hence, differences in drift rate may reflect the differences in the ability to extract relevant information from a perceptual scene (in this case, an array of concurrently presented sounds). In the following section, we will discuss this relationship in more detail.

### 4.2 N2ac amplitudes predict drift rate and accuracy

To date, little is known about the functional relevance of the N2ac component. The regression analysis conducted here revealed that N2ac amplitudes significantly predicted variations in accuracy as well as drift rate, while they were unrelated to mean RTs, threshold separation, or non-decision time. These findings add to the sparse literature that has so far investigated the N2ac component in different contexts (Gamble & Luck, 2011; Gamble & Woldorff, 2015; Gamble & Woldorff, 2015; Klatt et al., 2018b; Lewald & Getzmann, 2015; Lewald et al., 2016). In addition, to our best knowledge, this is the first study to show an N2ac component in a sample of older adults. Gamble and Luck (2011) originally proposed that the N2ac arises in order to resolve the competition between simultaneously present stimuli and reflects the attentional orienting towards a target. They further elucidated that this may be based on the biasing of neural coding towards the attended stimulus, as observed in the visual modality. In fact, the observed relationship of N2ac amplitudes and drift rate may support this line of reasoning: Drift rate conceptually reflects the quality of relevant information derived from sensory input that eventually drives the decision process (Ratcliff, Spieler, & McKoon, 2000). Hence, the better participants may be able to resolve competition between concurrent sounds by focusing on the target (i.e., N2ac amplitude), the better the quality of information that prompts participants to make a decision (i.e., drift rate; or in other words, the higher the rate of evidence accumulation in favor of a given response). In turn, it logically follows that the better or more consistently participants are able to focus their attention onto a relevant target sound (i.e., N2ac amplitude), the higher their overall accuracy.

Interestingly, in addition to the similar N2ac amplitudes for younger and older adults, we found no significant interactions between N2ac amplitudes and age, neither for accuracy nor for drift rate (cf. Table 1). This may suggest that the variances within age groups contributes more strongly to the observed relationship than the variance between age groups. However, the difficulties of interpreting a null effect, such as a missing interaction with age, need to be considered as a caveat here. Although regression lines in Figure 6 show a trend toward an interaction of N2ac amplitude and age group, the calculated BFs (cf. *3.4*) suggest the data to be insensitive to age group differences, providing no substantial evidence in favor of the null or alternative hypothesis. Nevertheless, one may raise the question, if lower N2ac amplitudes result in lower drift rates and decreased performance, why did older adults not show reduced N2ac amplitudes, given that they performed significantly worse than the younger adults? On the one hand, the well-pronounced N2ac component in older adults may, at least in part, have resulted from the recruitment of additional top-down resources to allow for more efficient target selection. This interpretation would be in line with the decline-compensation hypothesis (Cabeza, Anderson, Locantore, & McIntosh, 2002; for review, see Schneider, Pichora-Fuller, & Daneman, 2010), proposing that age-related declines in peripheral and central auditory processing are compensated for by increased allocation of cognitive resources. Increases in attentional focussing, however, might not be sufficient to completely compensate for the reduced performance of the older group. On the other hand, we cannot exclude that we simply failed to find a significant difference in N2ac amplitudes due to a lack of power, as the calculation of BFs provided no substantial evidence in favor of the null hypothesis.

### 4.3 Is post-stimulus alpha power lateralization functionally relevant?

The present study also investigated alpha lateralization as a measure of attentional orienting within an auditory scene. Typically, alpha lateralization manifests in a bilateral decrease of alpha power, that is more pronounced over the contralateral hemisphere (relative to a target or a cue). This spatially specific modulation of oscillatory activity has been repeatedly associated with the top-down controlled voluntary allocation of attention (Foxe, Simpson, & Ahlfors, 1998; Haegens et al., 2011; Ikkai et al., 2016; Thut et al., 2006). Here, we replicated this consistently observed response in the alpha frequency band in a sample of younger and older participants who performed an auditory localization task, requiring them to indicate the location of a pre-defined target stimulus among three concordantly presented distractors. Our results suggested that older adults may in principle be able to recruit the same oscillatory mechanisms as younger adults when searching for a target among simultaneously present distractors (Klatt et al., 2018b). Although bayesian statistics were indecisive in whether the non-significant difference in alpha lateralization between age groups presents a true null effect, the preserved post-stimulus alpha lateralization corroborated a number of studies, showing intact alpha lateralization in older adults when anticipating an upcoming (lateralized) stimulus (Heideman et al., 2018; Leenders, Lozano-Soldevilla, Roberts, Jensen, & De Weerd, 2018; Tune et al., 2018). However, recent studies did not find alpha lateralization in older adults, although they were still able to perform the task as well as their younger counterparts (Hong et al., 2015; van der Waal, Farquhar, Fasotti, & Desain, 2017). This poses the question to what extent lateralized alpha dynamics are functionally relevant for behavior.

It is relatively undisputed that alpha power lateralization tracks the locus and timing of spatial attention (Bae & Luck, 2018; Foster et al., 2017; Samaha, Iemi, & Postle, 2017). In addition, a growing body of evidence supports the notion that the alpha rhythm as a correlate of spatial attention, so far predominantly investigated in the visual attention literature, analogously operates in different modalities (Haegens et al., 2011; Klatt et al., 2018a, 2018b; Thorpe, D’Zmura, & Srinivasan, 2012; Wöstmann et al., 2016, 2018). Yet, what remains a matter of debate is, (1) *how* alpha power lateralization aids selective spatial attention and (2) whether it reflects a necessary prerequisite for successful behavioral performance. Regarding the *how*, two prevailing views exist: The gating by inhibition theory, proposed by Jensen and Mazaheri (2010), suggested that the relative increase of alpha power over the ipsilateral hemisphere inhibits regions processing irrelevant information. Alternatively, it has been suggested that the relative decrease of alpha power over the contralateral hemisphere results in increased cortical excitability, allowing for enhanced processing of the targets (Noonan et al., 2016; Yamagishi et al., 2005). Both mechanisms are not necessarily mutually exclusive. In fact, Foster and Awh (2018) just recently pointed out that a lot of the empirical evidence is compatible with either the target enhancement or the distractor suppression account. Recent evidence suggested that both mechanisms might independently contribute to attentional orienting (Schneider et al., 2019). In line with those latter findings, Capilla, Schoffelen, Paterson, Thut, and Gross (2014) proposed distinct sources and behavioral correlates for the ipsilateral and contralateral portion of the alpha power signal.

Adressing the second question – *Does alpha lateralization reflect a necessary prerequisite for successful behavioral performance?* – a range of spatial-cueing studies has provided compelling evidence showing behavioral performance to be predicted by the degree of alpha lateralization (Haegens et al., 2011; Kelly et al., 2009; Thut et al., 2006). On the contrary, our findings question the notion that alpha power lateralization reflects a behaviorally relevant attentional mechanism: Surprisingly, we did not find any association between alpha lateralization and diffusion model parameters, mean RTs, or accuracy. This could be explained by the fact that the present study differed from the majority of previous studies in that it investigated alpha power modulations following stimulus presentation. That is, while alpha lateralization may in fact be necessary to successfully shift one’s attention *in anticipation of* an upcoming stimulus, it does not appear to be a required neural response in the attentional processing *following* the presentation of a multi-sound array. This is in line with the proposal previously made by van Ede et al. (2014), who similarly concluded that the relevance of attentional modulations might be “restricted to situations in which attention influences perception through anticipatory processes” (p. 139). However, in contrast to our results, these authors found that alpha lateralization was completely abolished during the processing of an ongoing tactile stimulus.

Alternatively, the lack of a relationship with behavioral performance may be due to the fact that we calculated a relative measure of alpha amplitudes, that is, the difference between ipsilateral and contralateral alpha power. In a cued somatosensory detection task, van Ede et al. (2012) found only contralateral alpha power amplitudes to be related to tactile detection performance, whereas fluctuations in the contralateral minus ipsilateral difference failed to predict performance. Similarly, other studies using a relative index of alpha power modulations did not find a strong relationship with behavioral performance (Limbach & Corballis, 2017; Tune et al., 2018). These findings, or rather null-findings, might strengthen the emerging view that both target enhancement (i.e., contralateral alpha power decrease) and distractor suppression (i.e., ipsilateral alpha power increase) differentially contribute to task performance (Schneider et al., 2019) and that this should be taken into account when analyzing the contribution of alpha power oscillations to behavior. Yet, it should be noted that there are studies that successfully demonstrated an effect of the relative strength of alpha lateralization on task performance (Haegens et al., 2011; Kelly et al., 2009), suggesting the reasons for those diverging results are likely to be more complex than just a methodological artifact. Also, it has to be noted that the respective BFs (below 1, but above 0.33) were rather indecisive; thus, although our data do not seem to support a significant relationship between alpha lateralization and behavioral performance, they cannot provide compelling evidence for a true null effect either.

Critically, one question remains unanswered: If alpha lateralization is not a necessary component of post-stimulus attentional processing in an auditory scene, what does it reflect? It might be that post-stimulus alpha lateralization is an “optional response” that may result in more effective target enhancement or distractor inhibition, when a specific strategy is applied. Hence, due to different strategies used by different participants, there might be no overall relationship between alpha lateralization and behavior when analyzed across all participants (Limbach & Corballis, 2017; Rihs, Michel, & Thut, 2009). Alternatively, as shown in a previous study using a very similar task design, auditory post-stimulus alpha lateralization might be more closely related to the spatial-specificity of the task (Klatt et al., 2018b). In the latter study, a lateralization of alpha power was only evident when participants were instructed to localize (instead of to simply detect) a target sound within a multi-sound array. Hence, we proposed that in post-stimulus attentional processing, the lateralization of alpha power indexes the access to a spatiotopic template that is used to generate a spatially-specific response (Klatt et al., 2018b). If alpha lateralization reflects such a process, one may argue that there should be no or a substantially reduced alpha lateralization in incorrect trials and thus, alpha lateralization should in fact be associated with behavioral performance. Such differences in ALI amplitudes for correct versus incorrect trials have in fact been previously reported (Tune et al., 2018; Wöstmann et al., 2016, 2018). The fact that we calculated alpha lateralization indices based on each participant’s mean alpha power in correctly answered trials may explain why we fail to capture such differences for a rather coarse, dichotic measure of behavioral performance such as accuracy.

## 5. Conclusion

In summary, fluctuations in N2ac amplitude predicted the rate of information accumulation (i.e., drift rate) as well as overall accuracy. We conclude that the N2ac component reflected the participants’ ability to resolve competition between co-occurring sounds by focusing on the target. This, in turn, determined the quality of the information accumulated during the decision-making process and thereby affected overall accuracy levels. In contrast, alpha lateralization was unrelated to behavioral performance, suggesting that successful attentional orienting within an auditory scene (as opposed to in anticipation of an upcoming target sound), does not rely on alpha lateralization. Our findings strengthen the proposal that alpha lateralization is not specific to the visual domain, but may reflect a supramodal attentional mechanism that generalizes to the auditory domain (Kerlin, Shahin, & Miller, 2010; Thorpe et al., 2012). Yet, we highlight that it is important to distinguish between cue-related, anticipatory modulations of alpha power and post-stimulus alpha power lateralization.

## Funding

This work was supported by the German Federal Ministry of Education and Research in the framework of the TRAIN-STIM project (grant number 01GQ1424E).

## Acknowledgements

The authors are grateful to David Schmude, Jonas Heyermann, Stefan Weber, and Michael-Christian Schlüter for their help in running the experiments, to Peter Dillmann and Tobias Blanke for preparing software and parts of the electronic equipment, and to two anonymous reviewers for valuable comments on a previous version of this manuscript.

## Declarations of interest

None.

## References

Alain, C., & McDonald, K. L. (2007). Age-related differences in neuromagnetic brain activity underlying concurrent sound perception. Journal of Neuroscience, 27(6), 1308–1314. http://doi.org/10.1523/JNEUROSCI.5433-06.2007

Bae, G.-Y., & Luck, S. J. (2018). Dissociable Decoding of Spatial Attention and Working Memory from EEG Oscillations and Sustained Potentials. The Journal of Neuroscience, 38(2), 409–422. http://doi.org/10.1523/JNEUROSCI.2860-17.2017

Bronkhorst, A. W. (2015). The cocktail-party problem revisited: early processing and selection of multi-talker speech. Attention, Perception, and Psychophysics, 77(5), 1465–1487. http://doi.org/10.3758/s13414-015-0882-9

Cabeza, R., Anderson, N. D., Locantore, J. K., & McIntosh, A. R. (2002). Aging Gracefully: Compensatory Brain Activity. NeuroImage, 17, 1394–1402. http://doi.org/10.1006/nimg.2002.1280

Capilla, A., Schoffelen, J. M., Paterson, G., Thut, G., & Gross, J. (2014). Dissociated α-band modulations in the dorsal and ventral visual pathways in visuospatial attention and perception. Cerebral Cortex, 24(2), 550–561. http://doi.org/10.1093/cercor/bhs343

Cherry, C. E. (1953). Some Experiments on the Recognition of Speech, with One and with Two Ears. The Journal of the Acoustical Society of America, 25(5), 975–979. http://doi.org/10.1121/1.1907229

Cook, R. D., & Weisberg, S. (1982). Residuals and Influence in Regression. New York: Chapman and Hall. Retrieved from https://conservancy.umn.edu/handle/11299/37076

Craddock, M., Poliakoff, E., El-deredy, W., Klepousniotou, E., & Lloyd, D. M. (2017). Pre-stimulus alpha oscillations over somatosensory cortex predict tactile misperceptions. Neuropsychologia, 96(September 2016), 9–18. http://doi.org/10.1016/j.neuropsychologia.2016.12.030

Cribari-Neto, F., & Zeileis, A. (2010). Beta Regression in R. Journal of Statistical Software, 34(2), 1–24. http://doi.org/10.18637/jss.v034.i02

Delorme, A., & Makeig, S. (2004). EEGLAB: an open source toolbox for analysis of single-trial EEG dynamics including independent component analysis. Journal of Neuroscience Methods, 134(2004), 9–21. http://doi.org/10.1016/j.jneumeth.2003.10.009

Dienes, Z. (2014). Using Bayes to get the most out of non-significant results. Frontiers in Psychology, 5(July), 1–17. http://doi.org/10.3389/fpsyg.2014.00781

Eimer, M. (1996). The N2pc component as an indicator of attentional selectivity. Electroencephalography and Clinical Neurophysiology, 99(3), 225–234. http://doi.org/10.1016/S0921-884X(96)95711-2

Foster, J. J., & Awh, E. (2018). The role of alpha oscillations in spatial attention: Limited evidence for a suppression account. Current Opinion in Psychology, 29, 34–40. http://doi.org/10.1016/J.COPSYC.2018.11.001

Foster, J. J., Sutterer, D. W., Serences, J. T., Vogel, E. K., & Awh, E. (2017). Alpha-band oscillations enable spatially and temporally resolved tracking of covert spatial attention. Psychological Science, 28(7), 929–941. http://doi.org/10.1177/0956797617699167

Foxe, J. J., Simpson, G. V., & Ahlfors, S. P. (1998). Parieto-occipital ~10 Hz activity reflects anticipatory state of visual attention mechanisms. NeuroReport, 9(17), 3929–3933. http://doi.org/10.1097/00001756-199812010-00030

Gamble, M. L., & Luck, S. J. (2011). N2ac: An ERP Component Associated with the Focusing of Attention Within an Auditory Scene. Psychophysiology, 48(8), 1057–1068. http://doi.org/10.1111/j.1469-8986.2010.01172.x

Gamble, M. L., & Woldorff, M. G. (2015a). Rapid context-based identification of target sounds in an auditory scene. Journal of Cognitive Neuroscience, 27(9), 1675–1684. http://doi.org/10.1162/jocn_a_00814

Gamble, M. L., & Woldorff, M. G. (2015b). The temporal cascade of neural processes underlying target detection and attentional processing during auditory search. Cerebral Cortex, 25(9), 2456–2465. http://doi.org/10.1093/cercor/bhu047

Gould, I. C., Rushworth, M. F., & Nobre, A. C. (2011). Indexing the graded allocation of visuospatial attention using anticipatory alpha oscillations. Journal of Neurophysiology, 105, 1318–1326. http://doi.org/10.1152/jn.00653.2010.

Haegens, S., Handel, B. F., & Jensen, O. (2011). Top-Down Controlled Alpha Band Activity in Somatosensory Areas Determines Behavioral Performance in a Discrimination Task. Journal of Neuroscience, 31(14), 5197–5204. http://doi.org/10.1523/JNEUROSCI.5199-10.2011

Haegens, S., Luther, L., & Jensen, O. (2012). Somatosensory anticipatory alpha activity increases to suppress distracting input. Journal of Cognitive Neuroscience, 24(3), 677–685. http://doi.org/10.1162/jocn_a_00164

Händel, B. F., Haarmeier, T., & Jensen, O. (2011). Alpha oscillations correlate with the successful inhibition of unattended stimuli. Journal of Cognitive Neuroscience, 23(9), 2494–2502. http://doi.org/10.1162/jocn.2010.21557

Hanenberg, C., Getzmann, S., & Lewald, J. (2019). Transcranial direct current stimulation of posterior temporal cortex modulates electrophysiological correlates of auditory selective spatial attention in posterior parietal cortex. Neuropsychologia, 131(August 2018), 160–170. http://doi.org/10.1016/j.neuropsychologia.2019.05.023

Hansen, J. C., & Hillyard, S. A. (1980). Endogeneous brain potentials associated with selective auditory attention. Electroencephalography and Clinical Neurophysiology, 49(3-4), 277–290. http://doi.org/10.1016/0013-4694(80)90222-9

Heideman, S. G., Rohenkohl, G., Chauvin, J. J., Palmer, C. E., van Ede, F., & Nobre, A. C. (2018). Anticipatory neural dynamics of spatial-temporal orienting of attention in younger and older adults. NeuroImage, 178(March), 46–56. http://doi.org/10.1016/j.neuroimage.2018.05.002

Hentschke, H., & Stüttgen, M. C. (2011). Computation of measures of effect size for neuroscience data sets. European Journal of Neuroscience, 34(July), 1887–1894. http://doi.org/10.1111/j.1460-9568.2011.07902.x

Holm, S. (1979). A Simple Sequentially Rejective Multiple Test Procedure. Scandinavian Journal of Statistics, 6(2), 65–70.

Hong, X., Sun, J., Bengson, J. J., Mangun, G. R., & Tong, S. (2015). Normal aging selectively diminishes alpha lateralization in visual spatial attention. NeuroImage, 106, 353–363. http://doi.org/10.1016/j.neuroimage.2014.11.019

Ikkai, A., Dandekar, S., & Curtis, C. E. (2016). Lateralization in alpha-band oscillations predicts the locus and spatial distribution of attention. PloS One, 11(5), e0154796. http://doi.org/10.1371/journal.pone.0154796

Jensen, O., & Mazaheri, A. (2010). Shaping functional architecture by oscillatory alpha activity: gating by inhibition. Frontiers in Human Neuroscience, 4, 186. http://doi.org/10.3389/fnhum.2010.00186

Kelly, S. P., Gomez-Ramirez, M., & Foxe, J. J. (2009). The strength of anticipatory spatial biasing predicts target discrimination at attended locations: A high-density EEG study. European Journal of Neuroscience, 30(11), 2224–2234. http://doi.org/10.1111/j.1460-9568.2009.06980.x

Kelly, S. P., Lalor, E. C., Reilly, R. B., & Foxe, J. J. (2006). Increases in Alpha Oscillatory Power Reflect an Active Retinotopic Mechanism for Distracter Suppression During Sustained Visuospatial Attention. Journal of Neurophysiology, 95(6), 3844–3851. http://doi.org/10.1152/jn.01234.2005

Kerlin, J. R., Shahin, A. J., & Miller, L. M. (2010). Attention gain control of ongoing cortical speech representations in a “cocktail party.” Journal of Neuroscience, 30(2), 620–628. http://doi.org/10.1523/JNEUROSCI.3631-09.2010.

Klatt, L. I., Getzmann, S., Wascher, E., & Schneider, D. (2018a). Searching for auditory targets in external space and in working memory: Electrophysiological mechanisms underlying perceptual and retroactive spatial attention. Behavioural Brain Research, 353, 98–107. http://doi.org/10.1016/j.bbr.2018.06.022

Klatt, L. I., Getzmann, S., Wascher, E., & Schneider, D. (2018b). The contribution of selective spatial attention to sound detection and sound localization: Evidence from event-related potentials and lateralized alpha oscillations. Biological Psychology, 138, 133–145. http://doi.org/10.1016/j.biopsycho.2018.08.019

Krekelberg, B. (2019). bayesFactor.

Leenders, M. P., Lozano-Soldevilla, D., Roberts, M. J., Jensen, O., & De Weerd, P. (2018) Diminished alpha lateralization during working memory but not during attentional cueing in older adults. Cerebral Cortex, 28(1), 21–32. http://doi.org/10.1093/cercor/bhw345

Lewald, J. (2016). Modulation of human auditory spatial scene analysis by transcranial direct current stimulation. Neuropsychologia, 84, 282–293. http://doi.org/10.1016/j.neuropsychologia.2016.01.030

Lewald, J. (2019). Bihemispheric anodal transcranial direct-current stimulation over temporal cortex enhances auditory selective spatial attention. Experimental Brain Research, (2018). http://doi.org/10.1007/s00221-019-05525-y

Lewald, J., & Getzmann, S. (2015). Electrophysiological correlates of cocktail-party listening. Behavioural Brain Research, 292, 157–166. http://doi.org/10.1016/j.bbr.2015.06.025

Lewald, J., Hanenberg, C., & Getzmann, S. (2016). Brain correlates of the orientation of auditory spatial attention onto speaker location in a “cocktail-party” situation. Psychophysiology, 53, 1484–1495. http://doi.org/10.1111/psyp.12692

Liang, F., Paulo, R., Molina, G., Clyde, M. A., & Berger, J. O. (2008). Mixtures of g priors for Bayesian variable selection. Journal of the American Statistical Association, 103(481), 410–423. http://doi.org/10.1198/016214507000001337

Lilliefors, H. W. (1967). On the Kolmogorov-Smirnov Test for Normality with Mean and Variance Unknown Hubert W. Lilliefors. Journal of the American Statistical Association, 62(318), 399–402.

Limbach, K., & Corballis, P. M. (2017). Alpha-power modulation reflects the balancing of task requirements in a selective attention task. Psychophysiology, 54(2), 224–234. http://doi.org/10.1111/psyp.12774

Luck, S. J. (2014). An introduction to the event-related potential technique (2nd ed.). MIT Press.

Luck, S. J., & Hillyard, S. A. (1994). Spatial Filtering During Visual Search: Evidence from Human Electrophysiology. Journal of Experimental Psychology: Human Perception and Performance, 20(5), 1000–1014.

Lüdecke, D. (2018). ggeffects: Tidy Data Frames of Marginal Effects from Regression Models. Journal of Open Source Software, 3(26), 772. http://doi.org/10.21105/joss.00772

Mathworks. (2019). Statistics and Machine Learning Toolbox ™ User’s Guide (R 2019b).

Mognon, A., Jovicich, J., Bruzzone, L., & Buiatti, M. (2011). ADJUST: An automatic EEG artifact detector based on the joint use of spatial and temporal features. Psychphysiology, 48, 229–240. http://doi.org/10.1111/j.1469-8986.2010.01061.x

Mok, R. M., Myers, N. E., Wallis, G., & Nobre, A. C. (2016). Behavioral and neural markers of flexible attention over working memory in aging. Cerebral Cortex, 26(4), 1831–1842. http://doi.org/10.1093/cercor/bhw011

Morey, R. D., & Rouder, J. N. (2018). BayesFactor: Computation of Bayes Factors for Common Designs. R package version 0.9.12-4.2. https://CRAN.R-project.org/package=BayesFactor.

Myers, N. E., Walther, L., Wallis, G., Stokes, M. G., & Nobre, A. C. (2015). Temporal dynamics of attention during encoding vs. maintenance of working memory: complementary views from event-related potentials and alpha-band oscillations. Journal of Cognitive Neuroscience, 27(3), 492–508. http://doi.org/10.1162/jocn_a_00727

Myerson, J., Hale, S., Wagstaff, D., Poon, L. W., & Smith, G. A. (1990). The information-loss model: A mathematical theory of age-related cognitive slowing. Psychological Review, 97(4), 475–487. http://doi.org/10.1037//0033-295x.97.4.475

Noonan, M. P., Adamian, N., Pike, X. A., Printzlau, F., Crittenden, B. M., & Stokes, M. G. (2016). Distinct Mechanisms for Distractor Suppression and Target Facilitation. The Journal of Neuroscience, 36(6), 1797–1807. http://doi.org/10.1523/JNEUROSCI.2133-15.2016

Nunez, M. D., Vandekerckhove, J., & Srinivasan, R. (2017). How attention influences perceptual decision making: Single-trial EEG correlates of drift-diffusion model parameters. Journal of Mathematical Psychology, 76, 117–130. http://doi.org/10.1016/j.jmp.2016.03.003

Oldfield, R. C. (1971). The assessment and analysis of handedness: the Edinburgh inventory. Neuropsychologia, 9, 97–113. Retrieved from http://www.ncbi.nlm.nih.gov/pubmed/5146491

Onton, J., & Makeig, S. (2006). Information-based modeling of event-related brain dynamics. In C. Neuper & W. Klimesch (Eds.), Progress in Brain Research (Vol. 159, pp. 99–120). Elsevier B.V. http://doi.org/10.1016/S0079-6123(06)59007-7

Philiastides, M. G., Ratcliff, R., & Sajda, P. (2006). Neural Representation of Task Difficulty and Decision Making during Perceptual Categorization: A Timing Diagram. The Journal of Neuroscience, 26(35), 8965–8975. http://doi.org/10.1523/JNEUROSCI.1655-06.2006

Pichora-Fuller, M. K., Alain, C., & Schneider, B. A. (2017). Older Adults at the Cocktail Party. In R. F. J. Middlebrooks, J. Simon, A. Popper (Ed.), The Auditory System at the Cocktail Party (Vol. 60, pp. 227–259). Cham: Springer. http://doi.org/10.1007/978-3-319-51662-2_9

Pituch, K. A., & Stevens, J. P. (2016). Applied Multivariate Statistics for the Social Sciences (6th edition). New York, NY: Routledge.

Quintana, D. S., & Williams, D. R. (2018). Bayesian alternatives for common null-hypothesis significance tests in psychiatry: a non-technical guide using JASP. BMC Psychiatry, 18(178), 1–8.

Ratcliff, R., & McKoon, G. (2008). The Diffusion Decision Model: Theory and Data for Two-Choice Decision Tasks. Neural Computation, 20(4), 873–922. http://doi.org/10.1162/neco.2008.12-06-420

Ratcliff, R., Philiastides, M. G., & Sajda, P. (2009). Quality of evidence for perceptual decision making is indexed by trial-to-trial variability of the EEG. Proceedings of the National Academy of Sciences, 106(16), 6539–6544. http://doi.org/10.1073/pnas.0812589106

Ratcliff, R., Spieler, D., & Mckoon, G. (2000). Explicitly modeling the effects of aging on response time. Psychonomic Bulletin and Review, 7(1), 1–25. http://doi.org/10.3758/BF03210723

Ratcliff, R., Thapar, A., & McKoon, G. (2001). The Effects of Aging on Reaction Time in a Signal Detection Task. Psychology and Aging, 16(2), 323–341. http://doi.org/10.1037/0882-7974.16.2.323

Ratcliff, R., Thapar, A., & McKoon, G. (2003). A diffusion model analysis of the effects of aging on brightness discrimination. Perception & Psychophysics, 65(4), 523–535. Retrieved from http://search.ebscohost.com/login.aspx?direct=true&db=psyh&AN=2004-14390-004&site=ehost-live&scope=site

Ratcliff, R., Thapar, A., & McKoon, G. (2011). The effects of aging and IQ on item and associative memory. Journal of Experimental Psychology. General, 140(3), 464–487. http://doi.org/10.1158/0008-5472.CAN-10-4002.BONE

Rihs, T. A., Michel, C. M., & Thut, G. (2007). Mechanisms of selective inhibition in visual spatial attention are indexed by α-band EEG synchronization. European Journal of Neuroscience, 25, 603–610. http://doi.org/10.1111/j.1460-9568.2007.05278.x

Rihs, T. A., Michel, C. M., & Thut, G. (2009). A bias for posterior α-band power suppression versus enhancement during shifting versus maintenance of spatial attention. NeuroImage, 44(1), 190–199. http://doi.org/10.1016/j.neuroimage.2008.08.022

Romei, V., Gross, J., & Thut, G. (2010). On the Role of Prestimulus Alpha Rhythms over Occipito-Parietal Areas in Visual Input Regulation: Correlation or Causation? Journal of Neuroscience, 30(25), 8692–8697. http://doi.org/10.1523/JNEUROSCI.0160-10.2010

Samaha, J., Iemi, L., & Postle, B. R. (2017). Prestimulus alpha-band power biases visual discrimination confidence, but not accuracy. Consciousness and Cognition, 54, 47–55. http://doi.org/10.1016/j.concog.2017.02.005

Sauseng, P., Klimesch, W., Stadler, W., Schabus, M., Doppelmayr, M., Hanslmayr, S., … Birbaumer, N. (2005). A shift of visual spatial attention is selectively associated with human EEG alpha activity. European Journal of Neuroscience, 22(11), 2917–2926. http://doi.org/10.1111/j.1460-9568.2005…Log

Schmiedek, F., Oberauer, K., Wilhelm, O., Süß, H. M., & Wittmann, W. W. (2007). Individual Differences in Components of Reaction Time Distributions and Their Relations to Working Memory and Intelligence. Journal of Experimental Psychology: General, 136(3), 414–429. http://doi.org/10.1037/0096-3445.136.3.414

Schneider, B. A., Pichora-Fuller, K., & Daneman, M. (2010). Effects of Senescent Changes in Audition and Cognition on Spoken Language Comprehension. In & R. R. F. S. Gordon-Salant, R.D. Frisna, A.N. Popper (Ed.), The Aging Auditory System (pp. 167–210). New York: Springer. http://doi.org/10.1055/s-2006-954865.

Schneider, D., Göddertz, A., Haase, H., Hickey, C., & Wascher, E. (2019). Hemispheric asymmetries in EEG alpha oscillations indicate active inhibition during attentional orienting within working. Behavioural Brain Research, 359, 38–46. http://doi.org/10.1016/j.bbr.2018.10.020

Schubert, A.-L., Hagemann, D., Voss, A., Schankin, A., & Bergmann, K. (2015). Decomposing the relationship between mental speed and mental abilities. Intelligence, 51, 28–46. http://doi.org/10.1016/j.intell.2015.05.002

Schubert, A.-L., Nunez, M. D., Hagemann, D., & Vandekerckhove, J. (2018a). Individual Differences in Cortical Processing Speed Predict Cognitive Abilities: a Model-Based Cognitive Neuroscience Account. Computational Brain & Behavior. http://doi.org/10.1007/s42113-018-0021-5

Schubert, A.-L., Nunez, M. D., Hagemann, D., & Vandekerckhove, J. (2018b). Individual Differences in Cortical Processing Speed Predict Cognitive Abilities: a Model-Based Cognitive Neuroscience Account. Computational Brain & Behavior, 1–21. http://doi.org/10.1007/s42113-018-0021-5

Shinn-Cunningham, B. G. (2017). Cortical and Sensory Causes of Individual Differences in Selective Attention Ability Among Listeners With Normal Hearing Thresholds. Journal of Speech, Language, and Hearing Research, 60(10), 2976–2988. http://doi.org/10.1044/2017_jslhr-h-17-0080

Snyder, J. S., & Alain, C. (2005). Age-related changes in neural activity associated with concurrent vowel segregation. Cognitive Brain Research, 24(3), 492–499. http://doi.org/10.1016/j.cogbrainres.2005.03.002

Spaniol, J., Madden, D. J., & Voss, A. (2006). A Diffusion Model Analysis of Adult Age Differences in Episodic and Semantic Long-Term Memory Retrieval. Journal of Experimental Psychology: Learning Memory and Cognition, 32(1), 101–117. http://doi.org/10.1037/0278-7393.32.1.101

Starns, J. J., & Ratcliff, R. (2010). The effects of aging on the speed-accuracy compromise: Boundary optimality in the diffusion model. Psychology of Aging, 25(1), 377–390. http://doi.org/10.1037/a0018022

Thapar, A., Ratcliff, R., & McKoon, G. (2003). A diffusion model analysis of the effects of aging on letter Discrimination. Psychology and Aging, 18(3), 415–429. http://doi.org/10.1037/0882-7974.19.2.278

Thorpe, S., D’Zmura, M., & Srinivasan, R. (2012). Lateralization of frequency-specific networks for covert spatial attention to auditory stimuli. Brain Topography, 25(1), 39–54. http://doi.org/1doi:10.1007/s10548-011-0186-x

Thut, G., Nietzel, A., Brandt, S. A., & Pascual-Leone, A. (2006). α-Band Electroencephalographic Activity over Occipital Cortex Indexes Visuospatial Attention Bias and Predicts Visual Target Detection. Journal of Neuroscience, 26(37), 9494–9502. http://doi.org/10.1523/JNEUROSCI.0875-06.2006

Tune, S., Wöstmann, M., & Obleser, J. (2018). Probing the limits of alpha power lateralization as a neural marker of selective attention in middle-aged and older listeners. European Journal of Neuroscience, 48(7), 2537–2550. http://doi.org/10.1101/267989

Turner, B. M., Rodriguez, C. A., Norcia, T. M., McClure, S. M., & Steyvers, M. (2016). Why more is better: Simultaneous modeling of EEG, fMRI, and behavioral data. NeuroImage, 128, 96–115. http://doi.org/10.1016/j.neuroimage.2015.12.030

Vaden, R. J., Hutcheson, N. L., McCollum, L. A., Kentros, J., & Visscher, K. M. (2012). Older adults, unlike younger adults, do not modulate alpha power to suppress irrelevant information. NeuroImage, 63(3), 1127–1133. http://doi.org/10.1016/j.neuroimage.2012.07.050

Van Der Lubbe, R. H. J., Bundt, C., & Abrahamse, E. L. (2014). Internal and external spatial attention examined with lateralized EEG power spectra. Brain Research, 1583, 179–192. http://doi.org/10.1016/j.brainres.2014.08.007

van der Waal, M., Farquhar, J., Fasotti, L., & Desain, P. (2017). Preserved and attenuated electrophysiological correlates of visual spatial attention in elderly subjects. Behavioural Brain Research, 317, 415–423. http://doi.org/10.1016/j.bbr.2016.09.052

van Dijk, H., Schoffelen, J.-M., Oostenveld, R., & Jensen, O. (2008). Prestimulus Oscillatory Activity in the Alpha Band Predicts Visual Discrimination Ability. Journal of Neuroscience, 28(8), 1816–1823. http://doi.org/10.1523/JNEUROSCI.1853-07.2008

van Driel, J., Gunseli, E., Meeter, M., & Olivers, C. N. L. (2017). Local and interregional alpha EEG dynamics dissociate between memory for search and memory for recognition. NeuroImage, 149, 114–128. http://doi.org/10.1016/j.neuroimage.2017.01.031

van Ede, F., Niklaus, M., & Nobre, A. C. (2017). Temporal expectations guide dynamic prioritization in visual working memory through attenuated α oscillations. The Journal of Neuroscience, 37(2), 437–445. http://doi.org/10.1523/JNEUROSCI.2272-16.2016

van Ede, F., Szebényi, S., & Maris, E. (2014). Attentional modulations of somatosensory alpha, beta and gamma oscillations dissociate between anticipation and stimulus processing. NeuroImage, 97, 134–141. http://doi.org/10.1016/j.neuroimage.2014.04.047

Voss, A., Nagler, M., & Lerche, V. (2013). Diffusion models in experimental psychology: A practical introduction. Experimental Psychology, 60(6), 385–402. http://doi.org/10.1027/1618-3169/a000218

Voss, A., & Voss, J. (2007). Fast-dm: A free program for efficient diffusion model analysis. Behavior Research Methods, 39(4), 767–775. http://doi.org/10.3758/BF03192967

Voss, A., Voss, J., & Lerche, V. (2015). Assessing cognitive processes with diffusion model analyses: A tutorial based on fast-dm-30. Frontiers in Psychology, 6, 336. http://doi.org/10.3389/fpsyg.2015.00336

Wagenmakers, E. J., Marsman, M., Jamil, T., Ly, A., Verhagen, J., Love, J., … Morey, R. D. (2018). Bayesian inference for psychology. Part I: Theoretical advantages and practical ramifications. Psychonomic Bulletin and Review, 25(1), 35–57. http://doi.org/10.3758/s13423-017-1343-3

Wildegger, T., van Ede, F., Woolrich, M., Gillebert, C. R., & Nobre, A. C. (2017). Preparatory α-band oscillations reflect spatial gating independently of predictions regarding target identity. Journal of Neurophysiology, 117(3), 1385–1394. http://doi.org/10.1152/jn.00856.2016

Worden, M. S., Foxe, J. J., Wang, N., & Simpson, G. V. (2000). Anticipatory biasing of visuospatial attention indexed by retinotopically specific α-band electroencephalography increases over occipital cortex. Journal of Neuroscience, 20(6), RC63. http://doi.org/10.1523/JNEUROSCI.20-06-j0002.2000

Wöstmann, M., Herrmann, B., Maess, B., & Obleser, J. (2016). Spatiotemporal dynamics of auditory attention synchronize with speech. Proceedings of the National Academy of Sciences of the United States of America, 113(14), 1523357113-. http://doi.org/10.1073/pnas.1523357113

Wöstmann, M., Vosskuhl, J., Obleser, J., & Herrmann, C. S. (2018). Opposite effects of lateralised transcranial alpha versus gamma stimulation on auditory spatial attention. Brain Stimulation, 11(4), 752–758. http://doi.org/10.1016/j.brs.2018.04.006

Yamagishi, N., Goda, N., Callan, D. E., Anderson, S. J., & Kawato, M. (2005). Attentional shifts towards an expected visual target alter the level of alpha-band oscillatory activity in the human calcarine cortex. Cognitive Brain Research, 25(3), 799–809. http://doi.org/10.1016/j.cogbrainres.2005.09.006

Zündorf, I. C., Karnath, H. O., & Lewald, J. (2011). Male advantage in sound localization at cocktail parties. Cortex, 47(6), 741–749. http://doi.org/10.1016/j.cortex.2010.08.002

Zündorf, I. C., Karnath, H. O., & Lewald, J. (2014). The effect of brain lesions on sound localization in complex acoustic environments. Brain, 137(5), 1410–1418. http://doi.org/10.1093/brain/awu044

Zündorf, I. C., Lewald, J., & Karnath, H. O. (2013). Neural Correlates of Sound Localization in Complex Acoustic Environments. PLoS ONE, 8(5), e63259. http://doi.org/10.1371/journal.pone.0064259

